# scReadSim: a single-cell RNA-seq and ATAC-seq read simulator

**DOI:** 10.1101/2022.05.29.493924

**Authors:** Guanao Yan, Dongyuan Song, Jingyi Jessica Li

## Abstract

Benchmarking single-cell RNA-seq (scRNA-seq) and single-cell ATAC-seq (scATAC-seq) computational tools demands simulators to generate realistic sequencing reads. However, none of the few read simulators aim to mimic real data. To fill this gap, we introduce scReadSim, a single-cell RNA-seq and ATAC-seq read simulator that allows user-specified ground truths and generates synthetic sequencing reads (in FASTQ and BAM formats) by mimicking real data. At both read-sequence and read-count levels, scReadSim mimics real scRNA-seq and scATAC-seq data. Moreover, scReadSim provides ground truths, including unique molecular identifier counts for scRNA-seq and open chromatin regions for scATAC-seq. In particular, scReadSim allows users to design cell-type-specific ground-truth open chromatin regions for scATAC-seq data generation. In benchmark applications of scReadSim, we show that cell-ranger is a preferred scRNA-seq UMI deduplication tool, and HMMRATAC and MACS3 achieve top performance in scATAC-seq peak calling.

## Introduction

The development of single-cell sequencing technologies has enabled the characterization of genomic, epigenomic, and transcriptomic features of individual cells [1]. More than a thousand computational tools have been developed to analyze single-cell sequencing data [2], necessitating third-party benchmarking of computational tools. Realistic simulators are essential for fair benchmarking, because they can generate synthetic data that mimic real data and contain ground truths for evaluating computational tools.

Although many simulators have been developed for the two most popular single-cell sequencing technologies—single-cell RNA sequencing (scRNA-seq) and single-cell Assay for Transposase-Accessible Chromatin using sequencing (scATAC-seq), most of them do not generate sequencing reads. Instead, they only simulate read counts in genes [3–7] or genomic regions [8], and we refer to them as “count simulators.” (See [9] for a comprehensive review of scRNA-seq count simulators.) As such, these count simulators cannot be used to benchmark read-level bioinformatics tools that process reads stored in FASTQ or BAM files (Fig. S1). Exemplary read-level tools include unique molecular identifier (UMI) deduplication tools for scRNA-seq data [10–12] and peak-calling tools for scATAC-seq data [13–16]. Benchmarking these tools requires simulators that generate synthetic sequencing reads, which we refer to as “read simulators.”

For scRNA-seq, minnow [17] and Dropify [18] are the only two read simulators to our knowledge. Only minnow has a publicly available software package, which inputs a UMI count matrix and a gene annotation file but does not learn from real scRNA-seq reads. Moreover, minnow generates scRNA-seq reads only from annotated genes, so it cannot resemble real scRNA-seq data in intergenic regions that might be transcribed.

For scATAC-seq, SCAN-ATAC-Sim [19] is the only read simulator to our knowledge. However, similar to minnow, SCAN-ATAC-Sim does not learn from real scATAC-seq reads. Instead, it generates scATAC-seq reads from bulk ATAC-seq reads under simplistic assumptions: treating genomic regions (peaks called from bulk ATAC-seq reads) as independent, with a maximum of 2 reads allowed in each genomic region per cell, irrespective of the region’s length. Moreover, SCAN-ATAC-Sim does not provide synthetic read sequences with quality scores in a FASTQ or BAM file. Rather, it generates a BED file that only contains read start and end positions, thereby limiting direct benchmarking of read-level tools that typically require input in the form of FASTQ or BAM files. Additionally, we found that installing the minnow and SCAN-ATAC-Sim software packages in C++ was non-trivial due to the specific operating system and compiler requirements.

Motivated by the limitations of existing read simulators (a detailed comparison of scReadSim with the existing read simulators is in Table S1), we developed scReadSim as a scRNA-seq and scATAC-seq read simulator that generates realistic synthetic data by mimicking real data. scReadSim inputs a scRNA-seq or scATAC-seq BAM file and outputs synthetic reads in the FASTQ and BAM formats (Fig. 1a–b). We verified that scReadSim generates synthetic scRNA-seq and scATAC-seq data that resemble real data at both the read-sequence and read-count levels. Moreover, scReadSim provides ground truths, including unique molecular identifier (UMI) counts for scRNA-seq and open chromatin regions for scATAC-seq. In particular, scReadSim enables users to specify cell-type-specific open chromatin regions for scATAC-seq read generation, and it also allows users to vary the cell number and sequencing depth in generating synthetic data.

**Figure 1:**
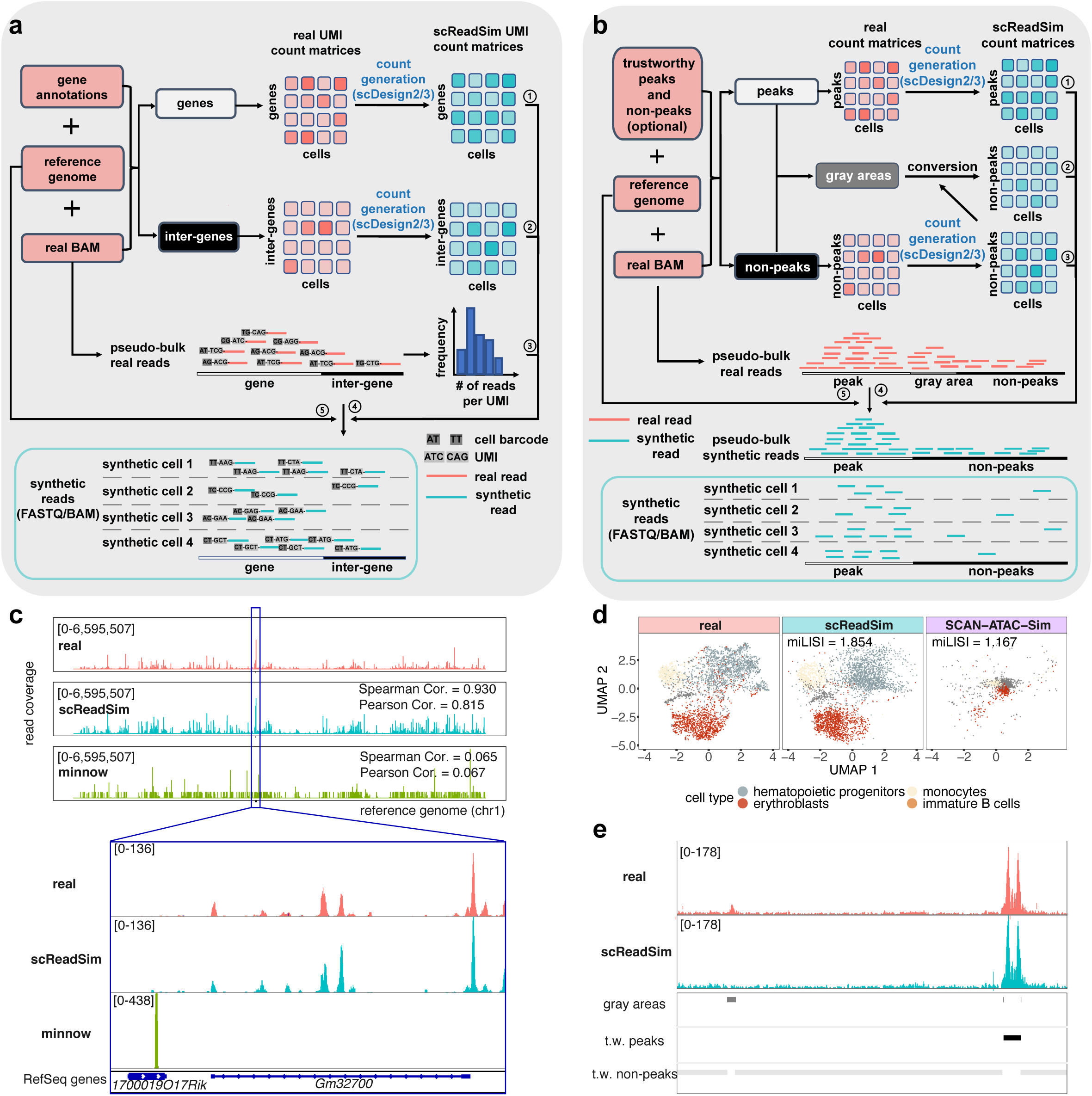
**a**, Workflow of scReadSim’s scRNA-seq read generation. The required input includes a real scRNA-seq dataset’s BAM file, the corresponding reference genome, and a gene annotation GTF file. Based on the input, scReadSim segregates the reference genome into genes and inter-genes (i.e., intergenic regions). Based on the genes and inter-genes, scReadSim summarizes scRNA-seq reads in the input BAM file into a gene-by-cell UMI count matrix and an inter-gene-by-cell UMI count matrix. Then scReadSim trains the count simulator scDesign2 [6] (if the cells belong to distinct clusters; otherwise, scDesign3 [21] can be used if the cells follow continuous trajectories) on the two UMI count matrices to generate the corresponding synthetic UMI count matrices (“ground truths” for benchmarking UMI deduplication tools). Last, scReadSim generates synthetic scRNA-seq reads based on the synthetic UMI count matrices (*0*1 and *0*2), the input BAM file (*0*3 and *0*4), and the reference genome (*0*5). The synthetic reads are outputted in FASTQ and BAM formats. **b**, Workflow of scReadSim’s scATAC-seq read generation. The input includes a real scATAC-seq dataset’s BAM file, the corresponding reference genome, and optionally, users’ trustworthy peaks and non-peaks in the input BAM file; if users do not input trustworthy peaks and non-peaks, scReadSim by default uses MACS3 [13] with stringent criteria to call trustworthy peaks (called at q-value 0.01) and non-peaks (complementary to the peaks called at q-value 0.1) from the input BAM file. Then scReadSim defines gray areas as the genomic regions complementary to the peaks and non-peaks, and scReadSim summarizes scATAC-seq reads in the input BAM file into a peak-by-cell count matrix and a non-peak-by-cell count matrix. Next, scReadSim trains the count simulator scDesign2 [6] (if the cells belong to distinct clusters; otherwise, scDesign3 [21] can be used if the cells follow continuous trajectories) on the two count matrices to generate the corresponding synthetic count matrices for the peaks and non-peaks. Further, scReadSim converts the gray areas into non-peaks (so that the peaks can be regarded as “ground-truth peaks”) and constructs a synthetic count matrix based on the gray areas’ lengths and the already-generated synthetic non-peak-by-cell count matrix. Last, scReadSim generates synthetic reads based on the three synthetic count matrices (*0*1, *0*2, and *0*3), the input BAM file (*0*4), and the reference genome (*0*5). The synthetic reads are outputted in FASTQ and BAM formats. **c**, scReadSim outperforms the existing scRNA-seq read simulator minnow in preserving the read coverage in a mouse 10x single-cell Multiome dataset (the RNA-seq modality only) [20]. With chromosome 1 of the reference genome divided into consecutive, non-overlapping 1,000-bp windows, the x-axis represents the windows’ positions along chromosome 1, and the y-axis indicates the number of the reads overlapping each window in each pseudo-bulk sample (real or synthetic, with all cells pooled). The track height is set to 6,595,507 for all three tracks. The Spearman correlation (Cor.) and Pearson Cor. measure the similarity of read coverage between real and synthetic data across the windows. Inset: a closer view of the region chr1:86,425,858–86,443,378 using the IGV genome browser [25]. Each coverage track displays the depth of the reads covered at each locus and indicates its track height 136 or 438 at the left corner. **d**, In UMAP visualizations, scReadSim outperforms the existing scATAC-seq read simulator SCAN-ATAC-Sim in mimicking real cells in the mouse 10x single-cell Multiome dataset (the ATAC-seq modality only) [20]. The miLISI measures the similarity of real and synthetic cells in the UMAP space: the miLISI value ranges between 1 and 2, with 2 indicating a perfect mixing of real and synthetic cells. **e**, scReadSim converts gray areas in real data to ground-truth non-peaks in synthetic data, maintaining trustworthy (t.w.) peaks and non-peaks in real data as ground-truth peaks and non-peaks, respectively, in synthetic data.

Among scReadSim’s myriad uses, we use two exemplary applications to highlight scReadSim’s utility as a benchmarking tool. First, for scRNA-seq, scReadSim mimics real data by first generating realistic UMI counts and then simulating reads. Hence, the synthetic UMI count matrix—an intermediate output of scReadSim—serves as the ground truth for benchmarking scRNA-seq UMI deduplication tools such as UMI-tools [10], cellranger [11], and Alevin [12], which all process reads (in a FASTQ file) into a UMI count matrix. Our benchmarking results indicate that cellranger is the preferred tool for its greater accuracy and higher computational efficiency. Second, for scATAC-seq, scReadSim provides ground-truth peaks and non-peaks by learning from user-specified trustworthy peaks and non-peaks or by calling trustworthy peaks and non-peaks from real data. The ground-truth peaks provided by scReadSim allow benchmarking peak-calling tools such as MACS3 [13], HOMER [14], HMMRATAC [15], and SEACR [16]. Our benchmarking results show that HMMRATAC and MACS3 are the top performers.

## Results

### scReadSim generates realistic synthetic scRNA-seq reads

To evaluate scReadSim’s ability to generate realistic scRNA-seq reads, we trained scReadSim on the RNA-seq modality of a mouse 10x single-cell Multiome dataset [20] and compared scReadSim’s synthetic data with the real data at two levels: UMI counts and read sequences. We also compared scReadSim with minnow, the only scRNA-seq read simulator with a publicly available software package.

At the UMI-count level, scReadSim uses realistic count simulators scDesign2 [6] and scDesign3 [21] to generate UMI counts (Fig. 1a; **Methods**), and both scDesign2 and scDesign3 have been verified to generate more realistic UMI counts compared with other count simulators. Hence, we only conducted a confirmation study by demonstrating that scReadSim’s synthetic UMI counts resemble real UMI counts in three aspects: (1) the distributions of six summary statistics, including four gene-level statistics (mean, variance, coefficient of variance, and zero proportion) and two cell-level statistics (zero proportion and cell library size) (Fig. S2a); (2) correlations among the top expressed genes (Fig. S2b); and (3) cells’ UMAP embeddings (Fig. S2c).

At the read-sequence level, we evaluated the resemblance of scReadSim’s synthetic reads to real reads in three aspects. First, scReadSim’s synthetic reads preserve the k-mer spectra, which describe the distribution of k-mers’ frequencies (i.e., numbers of occurrences) in reads. Fig. S3a shows similar k-mer spectra in synthetic reads and real reads. Second, scReadSim can introduce substitution errors to imitate those observed in real reads. Fig. S3b shows that scReadSim’s synthetic reads mimic real reads in terms of the substitution error rate per base call. Third, after read alignment, the pseudo-bulk read coverage of scReadSim’s synthetic reads mimics that of real reads (Spearman correlation = 0.930 in Fig. 1c; Fig. S3c). In contrast, the pseudo-bulk read coverage of minnow’s synthetic reads does not mimic real data (Spearman correlation = 0.065 Fig. 1c), an expected result as minnow does not learn from real scRNA-seq reads.

### scReadSim generates realistic synthetic scATAC-seq reads

For scATAC-seq, scReadSim provides ground-truth peaks and non-peaks for benchmarking purposes, by learning from user-specified or scReadSim-identified (under stringent criteria) peaks and non-peaks that are deemed trustworthy in real data (Fig. 1b; **Methods**). In synthetic scATAC-seq data generation, scReadSim first sets trustworthy peaks and non-peaks as ground-truth peaks and non-peaks, respectively. Then regarding the gray areas (i.e., regions complementary to trustworthy peaks and non-peaks), whose read coverages are between those of trustworthy peaks or non-peaks in real data, scReadSim converts them to ground-truth non-peaks (**Methods**).

We verified that scReadSim generates realistic synthetic scATAC-seq reads at two levels: read counts and read sequences, by mimicking two real datasets: the ATAC-seq modality of a mouse 10x single-cell Multiome dataset [20] and a sci-ATAC-seq dataset [22]. We also compared scReadSim with SCAN-ATAC-Sim, the only scATAC-seq read simulator available.

At the read-count level, we first confirmed that scReadSim’s synthetic peak-by-cell and non-peak-by-cell count matrices mimic the real count matrices in four aspects: (1) the distributions of six summary statistics, including four peak-level statistics (mean, variance, coefficient of variance, and zero proportion) and two cell-level statistics (zero proportion and cell library size) (Figs. S4a and S5a); (2) the distributions of normalized read counts (in Reads Per Kilobase Million (RPKM)) of peaks and non-peaks (Figs. S4b and S5b); (3) correlations among the top open peaks (Figs. S4c and S5c); and (4) cells’ UMAP embeddings (Figs. S4d and S5d). Second, we found that scReadSim outperforms SCAN-ATAC-Sim in mimicking the sci-ATAC-seq data in both the distributions of six summary statistics (Fig. S6a) and the cells’ UMAP embeddings (Figs. 1d and S6b), an expected result as SCAN-ATAC-Sim does not learn from real scATAC-seq data.

At the read-sequence level, we evaluated the resemblance of scReadSim’s synthetic scATAC-seq reads to real reads in five aspects. First, after read alignment, scReadSim’s synthetic read coverage mimics the real read coverage in the trustworthy peaks and non-peaks at the pseudo-bulk level (Figs. 1e and S7a); moreover, scReadSim’s synthetic read coverage in the ground-truth non-peaks converted from gray areas mimics the real read coverage in the nearby trustworthy non-peaks. We further verified that scReadSim preserves the cell-type-specific read coverage in real data (Fig. 2a). Second, scReadSim’s synthetic reads mimic real reads in terms of k-mer spectra (Figs. S7b and S8a). Third, scReadSim’s synthetic reads preserve the fragment size distribution of real reads (Figs. 2b Top and S8b); it is known that the fragment size distribution results from the pattern of chromatin accessibility and thus meaningful [23]. Fourth, scReadSim’s synthetic reads maintain the substitution error rate per base call in real reads (Figs. 2b Middle and S8c). Fifth, MACS3, a state-of-the-art peak calling tool, achieves consistent peak calling results from scReadSim’s synthetic reads and real reads (Figs. 2b Bottom, S7c and S8d–e; a reasonable result is that the consistency is higher for the ATAC-seq modality in the mouse 10x single-cell Multiome dataset, which is less sparse than the sci-ATAC-seq dataset).

**Figure 2:**
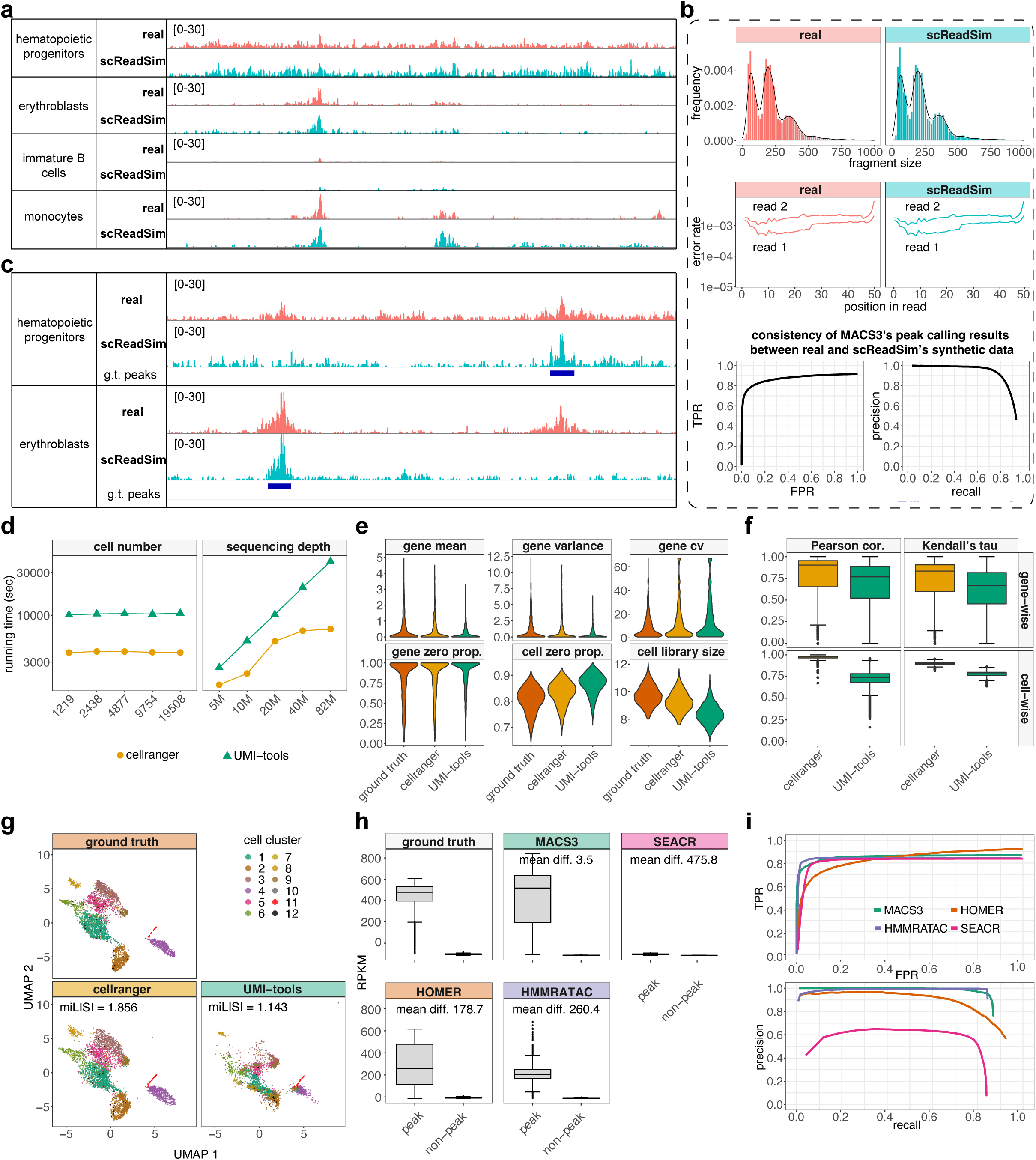
**a–b**, scReadSim’s synthetic scATAC-seq data mimic the real sci-ATAC-seq dataset [22] in terms of the cell-type-specific read coverage (**a**), the fragment-size distribution (**b** Top), substitution error rate per base call within a read (**b** Middle), and the peaks called by MACS3 at the pseudo-bulk level (**b** Bottom; **Methods**). **c**, scReadSim enables user-designed, cell-type-specific ground-truth (g.t.) peaks for scATAC-seq read generation. **d–g**, Benchmark of UMI deduplication tools using scReadSim’s synthetic scRNA-seq reads in four aspects: **d**, Time usage of deduplication tools on synthetic datasets with varying cell numbers (at a fixed sequencing depth) or varying sequencing depths (at a fixed cell number). The y-axis indicates the time lapse (in seconds), and the x-axis shows the number of synthetic cells (left) or the total number of UMIs (sequencing depth, right). **e**, Distributions of summary statistics of the UMI count matrices (ground truth, cellranger’s output, and UMI-tools’ output) at the gene level (mean, variance, coefficient of variance (cv), and zero proportion) and the cell level (zero proportion and library size). **f**, Cell-wise and gene-wise correlations (Pearson correlation and Kendell’s tau) between the ground-truth UMI count matrix and each deduplication tool’s output UMI count matrix. **g**, UMAP visualizations of the UMI count matrices. Cells are colored by the cell clusters outputted by scReadSim (the clusters are from the real data used to train scReadSim). The miLISI measures the similarity of real and synthetic cells in the UMAP space: the miLISI value ranges between 1 and 2, with 2 indicating a perfect mixing of real and synthetic cells. **h–i**, Benchmark of peak calling tools using scReadSim’s synthetic scATAC-seq reads through in two aspects: **h**, Distributions of RPKM values of peak and non-peak regions in the ground truth (specified in scReadSim) and each tool’s peak-calling result. Mean differences (mean diff.) of peak regions’ RPKM are calculated between ground truth and each method. **i**, true positive rate (TPR) vs. false positive rate (FPR) curves (top) and precision vs. recall curves (bottom) using user-designed open chromatin regions as the ground-truth peaks. A called peak is considered true if it overlaps at least 225 bp of a user-designed open chromatin region, where 225 bp is half of the shortest open chromatin region.

Beyond mimicking real data, scReadSim also allows users to design ground-truth peaks and non-peaks for synthetic scATAC-seq data generation at the cell-type level (Fig. 2c) or even arbitrarily (Fig. S9). We show that scReadSim’s synthetic scATAC-seq data with user-designed ground-truth peaks and non-peaks preserve the real read-sequence characteristics and mimic the trustworthy peaks and non-peaks in real data (Fig. S10). In summary, scReadSim can provide reliable ground truths for benchmarking computational tools that process scATAC-seq reads.

### scReadSim allows varying cell numbers and sequencing depths

scReadSim flexibly allows users to vary cell numbers and sequencing depths to generate synthetic data. For UMI-based scRNA-seq data, scReadSim defines the baseline sequencing depth as the number of UMIs in the input real data; for non-UMI-based scRNA-seq data and scATAC-seq data, scReadSim defines the baseline sequencing depth as the number of sequencing reads (with duplicated reads resulted from Polymerase Chain Reaction (PCR) removed) in the input real data. For demonstration purposes, we deployed scReadSim to the RNA-seq modality of a mouse 10x single-cell Multiome dataset [20] and varied the cell number or the sequencing depth (Fig. S11). Fig. S11a shows the synthetic datasets scReadSim generated with a fixed sequencing depth but increasing cell numbers 1,219 (0.25*×*), 2,438 (0.5*×*), 4,877 (1*×*, the original cell number), 9,754 (2*×*), and 19,508 (4*×*). For a fair visual comparison, we downsampled every synthetic dataset to 1,219 cells (0.25*×*) in Fig. S11a and observed that the read coverage of 1,219 synthetic cells decreases as the synthetic cell number increases, an expected phenomenon reflecting the trade-off between the cell number and the per-cell sequencing depth given a fixed sequencing depth.

Fig. S11b shows to the synthetic datasets scReadSim generated with a fixed cell number but increasing sequencing depths 1.2*M* (0.25*×*), 2.5*M* (0.5*×*), 5.0*M* (1*×*, the original sequencing depth), 10.1*M* (2*×*), and 20.1*M* (4*×*). As expected, the read coverage increases as the sequencing depth increases.

Note that although scReadSim allows users to specify the sequencing depth arbitrarily, users must consider the biological limits so that the specified sequencing depth is reasonable (e.g., the total number of mRNA transcripts is the upper bound on the number of UMIs).

### Application 1: Benchmarking UMI deduplication tools

For UMI-based scRNA-seq data, UMI deduplication tools were developed to quantify gene expression levels from scRNA-seq reads, some of which may come from the same RNA molecule. UMI deduplication tools input scRNA-seq reads (containing UMIs and cell barcodes) and output a gene-by-cell UMI count matrix. scReadSim can be used for benchmarking UMI deduplication tools because it provides the ground-truth UMI count matrix in its synthetic scRNA-seq read generation process. Hence, we deployed scReadSim to the mouse 10x single-cell Multiome dataset [20] (the RNA-seq modality only) and used the synthetic reads to benchmark popular UMI deduplication tools: cellranger [11], UMI-tools [10], and Alevin [12]. Since Alevin is a transcript-level deduplication tool but scReadSim generates gene-level scRNA-seq reads, we focus on cellranger and UMI-tools in the following result comparison (Fig. 2d–g). We only include Alevin’s result in Figs. S12 and S13.

Our benchmark study shows that cellranger achieves better performance than UMI-tools in three aspects. First, cellranger is computationally more efficient than UMI-tools on all synthetic datasets (with varying cell numbers and sequencing depths) (Fig. 2d). As expected, cellranger and UMI-tools both take longer to run when the sequencing depth increases, and both methods’ running time is unaffected by the cell number when the sequencing depth is fixed.

Second, the UMI count matrix output by cellranger agrees better with the ground-truth UMI count matrix in terms of (1) the distributions of six summary statistics, including four gene-level statistics (mean, variance, coefficient of variance, and zero proportion) and two cell-level statistics (zero proportion and cell library size) (Fig. 2e); and (2) Pearson correlation and Kendall’s tau, both gene-wise (the correlation across cells per gene) and cell-wise (the correlation across genes per cell) (Fig. 2f).

Third, UMAP visualization shows that, compared to UMI-tools, cellranger outputs a UMI count matrix that is more similar to the ground-truth UMI count matrix (evidenced by higher values (1.856) of miLISI (mean integration Local Inverse Simpson’s Index)) and better preserves the separation among cell clusters (Fig. 2g).

### Application 2: Benchmarking scATAC-seq peak-calling tools

For benchmarking scATAC-seq peak-calling tools, scReadSim provides ground-truth peaks and non-peaks by learning from user-specified trustworthy peaks and non-peaks or calling trustworthy peaks and non-peaks from real data.

Here we used scReadSim’s synthetic scATAC-seq data (pseudo-bulk), which contain designed ground-truth peaks and non-peaks (**Methods**) and mimic the sci-ATAC-seq dataset [22], to benchmark four popular peak-calling tools: (1) MACS3, designed for identifying transcript factor binding sites from ChIP-seq data; (2) SEACR, designed for CUT&RUN data; (3) HOMER, designed for ChIP-seq data but also widely used for ATAC-seq data; and (4) HMMRATAC, designed for ATAC-seq data.

Our benchmark results indicate that MACS3 and HMMRATAC have the best overall performance among the four peak-calling tools. In particular, the peaks called by MACS3 best resemble the ground-truth peaks in terms of read coverage, while HMMRATAC achieves the best balance between precision and recall.

First, we plot the RPKM distributions in ground-truth peaks and non-peaks, as well as in the identified peaks and non-peaks by each peak-calling tool (Fig. 2h). The distributions indicate that MACS3 identifies the peaks that are most similar to the ground-truth peaks, followed by HMMRATAC, HOMER, and SEACR. Specifically, the peaks called by HMMRATAC and SEACR are much longer than those by MACS3 and HOMER: the average peak lengths are 1543.0 bp and 1746.9 bp for HMMRATAC and SEACR, in contrast to 490.4 bp and 431.6 bp for MACS3 and HOMER. Possibly due to the design for CUT&RUN data, SEACR identifies the genomic regions with non-zero read coverages as peaks. However, the sci-ATAC-seq pseudo-bulk data used here contain many trustworthy non-peaks with non-zero read coverages, so most of the peaks called by SEACR contain so few reads that these peaks’ RPKMs are close to 0.

Second, all four tools detect most of the ground-truth peaks, under the assumption that a ground-truth peak is considered as correctly identified if a minimum length of the peak (i.e., half of the length of the shortest peak) is overlapped by an identified peak (Figs. 2i Left, S14–S15). Under the default settings of the four tools, the recall rates (i.e., the percentage of ground-truth peaks identified correctly) are all high: 86.8% for MACS3, 83.9% for SEACR, 92.4% for HOMER, and 84.4% for HMMRATAC.

Third, HMMRATAC achieves the highest precision rate of 90.3% (i.e., the percentage of identified peaks that are correct), followed by 76.5% for MACS3, 56.3% for HOMER, and 7.8% for SEACR. SEACR’s low precision rate is possibly due to its design, leading to a vast number of identified peaks (34,834 peaks vs. 3,678 peaks of MACS3, 5,314 peaks of HOMER, and 3,029 peaks of HMMRATAC) (Figs. 2i Right). Combining the precision and recall rates, the F1 scores indicate that HMMRATAC (87.3%) outperforms MACS3 (81.3%), HOMER (70.0%), and SEACR (14.3%).

### Summary

scReadSim, as a single-cell RNA-seq and single-cell ATAC-seq read simulator, generates realistic synthetic reads by mimicking real reads while providing the ground truths, including UMI count matrix for scRNA-seq and user-designed open chromatin regions for scATAC-seq. For both modalities, scReaSim allows the cell number and the sequencing depth to vary. Although the current version of scReadSim uses scDesign2 [6] as the internal count simulator (assuming that cells are from discrete cell types), scReadSim is easily adaptive to the new count simulator scDesign3 [21] that allows cells to come from continuous trajectories and have spatial locations. Moreover, scReadSim can be used for evaluating the robustness of read-level bioinformatics tools because scReadSim allows the generation of sequencing reads with varying levels of noise injected (e.g., read positions and coverages). As a future direction, we will generalize scReadSim to generate scRNA-seq short and long reads with ground-truth RNA isoform abundances, a functionality unavailable in existing scRNA-seq simulators.

## Data availability

- **10x Genomics**: The 10x Genomics single-cell multiome dataset includes ATAC and gene expression data from the embryonic mouse brain tissue [20]. The raw BAM files were downloaded from https://www.10xgenomics.com/resources/datasets/fresh-embryonic-e-18-mouse-brain-5-k-1-standard-2-0-0. The reference genome file (assembly version GRCm38.p4 release M10) was downloaded from GENCODE (https://www.gencodegenes.org/mouse/ release_M10.html).
- **sci-ATAC-seq**: The sci-ATAC-seq dataset measures the chromatin accessibility in 17 samples spanning 13 tissues in 8-week old mice [22]. The raw BAM file was downloaded from https://atlas.gs.washington.edu/mouse-atac/. Data from the tissue “Bone Marrow 62016” were used for analysis. The reference genome file (assembly version NCBIM37 release M1) was downloaded from GENCODE (https://www.gencodegenes.org/mouse/ release_M1.html).

## Code availability

The scReadSim Python package is available at https://github.com/JSB-UCLA/scReadSim. The comprehensive tutorials are available at http://screadsim.readthedocs.io/. In the tutorials, we described the input and output formats, model parameters, and exemplary datasets for each functionality of scReadSim. The source code for reproducing the results are available at: https://github.com/Dominic7227/scReadSim_source [24].

To run scReadSim, the following dependent software packages are needed.

- **MACS3**: Version 3.0.0a7 (https://github.com/macs3-project/MACS).
- **samtools**: Version 1.12 (with htslib 1.12) (http://www.htslib.org/).
- **bedtools**: Version 2.29.1 (https://bedtools.readthedocs.io/en/latest/).
- **seqtk**: Version 1.3-r117-dirty (https://github.com/lh3/seqtk).
- **bowtie2**: Version 2.3.4.1 (http://bowtie-bio.sourceforge.net/bowtie2/index.shtml).
- **Seurat**: R package Seurat Version 4.0.6.
- **fgbio**: Version 2.0.1 (http://fulcrumgenomics.github.io/fgbio/).

In addition, the following software packages were used for the analyses in this work.

### Existing read simulators

- **minnow**: Version 0.1.0 (https://github.com/COMBINE-lab/minnow)
- **SCAN-ATAC-Sim**: http://scan-atac-sim.gersteinlab.org/

### Dimension reduction methods

- **UMAP**: Function umap() from R package umap Version 0.2.7.0.
- **PCA**: Function irlba() from R package irlba Version 2.3.5.

### UMI deduplication tools

- **UMI-tools**: Version 1.1.2 (https://github.com/CGATOxford/UMI-tools).
- **Alevin**: Software Alevin (integrated into the software salmon Version 1.8.0) (https://salmon. readthedocs.io/en/latest/alevin.html).
- **cellranger**: Version 7.0.0 (https://support.10xgenomics.com/single-cell-gene-expression/ software/pipelines/latest/what-is-cell-ranger).

### Peak-calling tools

- **MACS3**: Version 3.0.0a7 (https://github.com/macs3-project/MACS).
- **SEACR**: Version 1.3 (https://github.com/FredHutch/SEACR).
- **HOMER**: Software HOMER Version 4.11 (http://homer.ucsd.edu/homer/index.html).
- **HMMRATAC**: Version 1.2.10 (https://github.com/LiuLabUB/HMMRATAC).

### Other tools

- **Jellyfish**: Version 2.3.0 (https://github.com/gmarcais/Jellyfish).
- **Intervene**: Python package Intervene Version 0.41.0 (https://intervene.readthedocs. io/en/latest/install.html).

## Competing interests

The authors declare no competing interests.

## Acknowledgements

The authors appreciate the comments and feedback from the members of the Junction of Statistics and Biology at UCLA (http://jsb.ucla.edu).

## Funding

This work was supported by the following grants: National Science Foundation DBI-1846216 and DMS-2113754, NIH/NIGMS R01GM120507 and R35GM140888, Johnson & Johnson WiSTEM2D Award, Sloan Research Fellowship, UCLA David Geffen School of Medicine W.M. Keck Foundation Junior Faculty Award, and Chan-Zuckerberg Initiative Single-Cell Biology Data Insights [Silicon Valley Community Foundation Grant Number: 2022-249355] (to J.J.L.).

## Supplementary Figures

**Figure S1:**
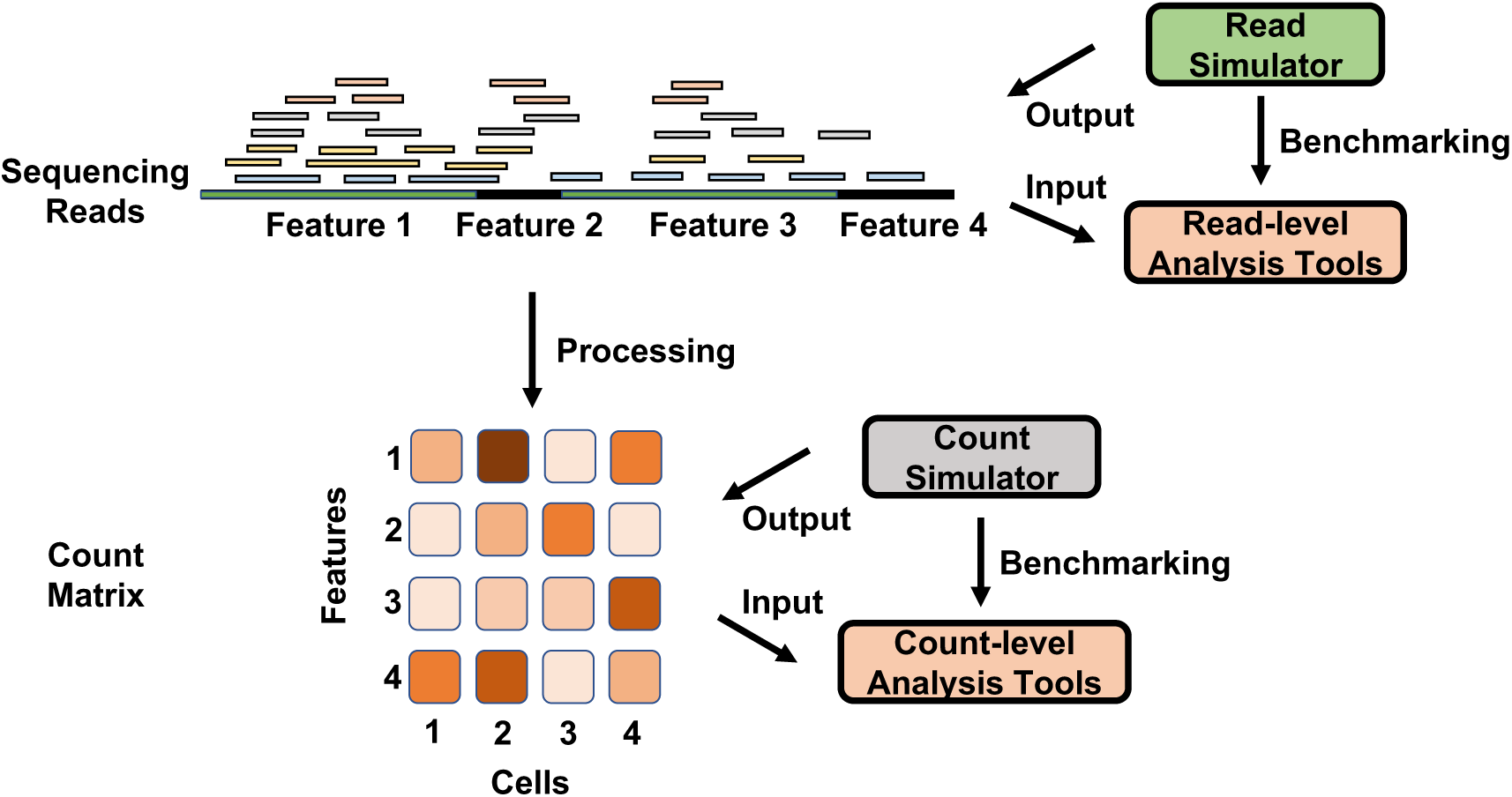
Relationships between single-cell sequencing data analysis tools and simulators. Read-level analysis tools process sequencing reads directly, while count-level analysis tools input a processed sequencing read count matrix. To benchmark count-level analysis tools, researchers need count simulators that generate synthetic sequencing read counts. However, count simulators cannot provide ground truths for benchmarking read-level analysis tools.

**Figure S2:**
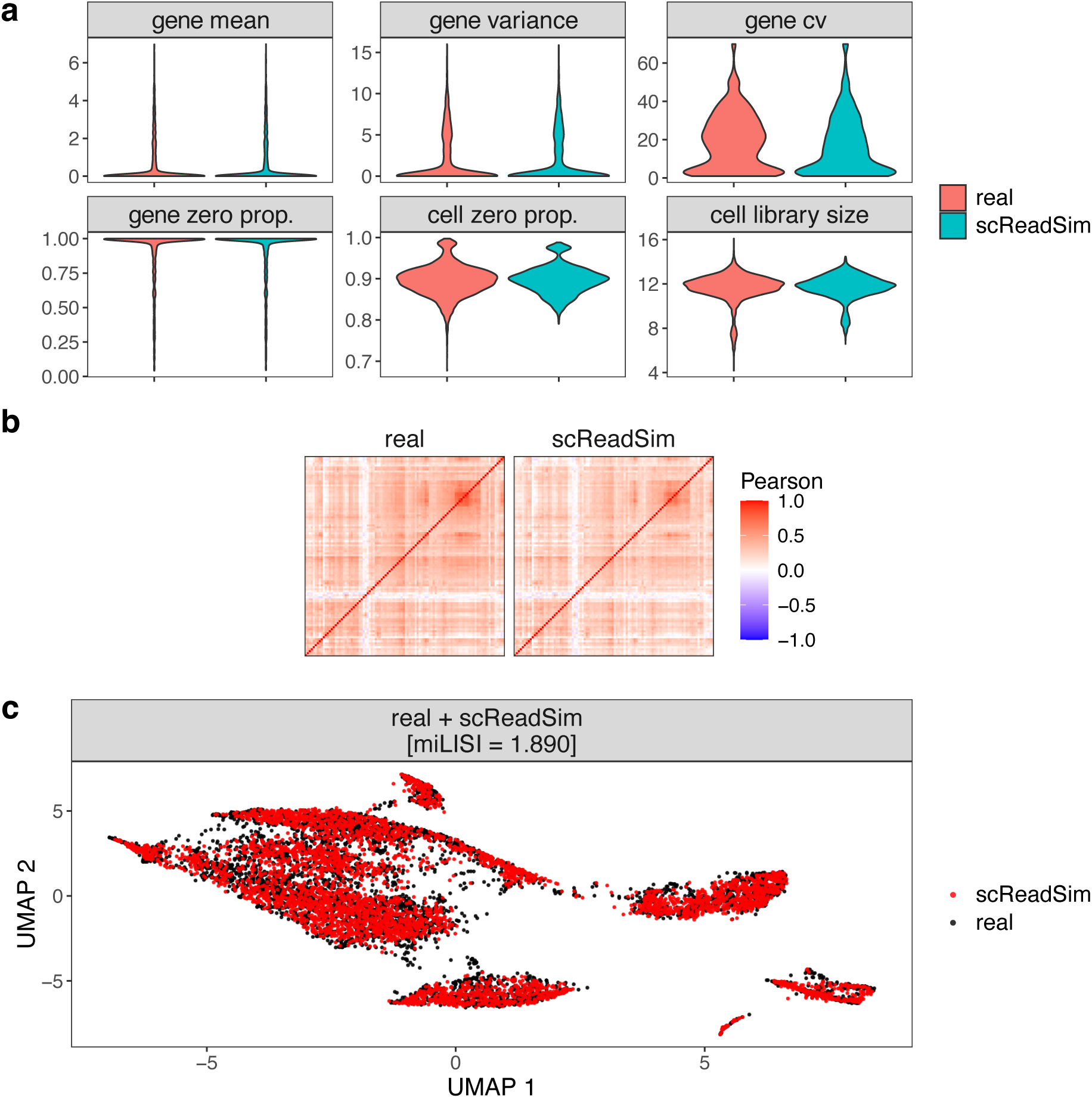
scReadSim’s synthetic data resemble real mouse 10x single-cell Multiome dataset (the RNA-seq modality only) [20] at the UMI-count level. **a**, Summary statistics of the UMI count matrix at the gene level (mean, variance, coefficient of variance (cv), and zero proportion) and the cell level (zero proportion and cell library size). **b**, Correlations among the 100 top-expressed genes in the synthetic and real count matrices. The top-expressed genes are defined based on the real count matrix. **c**, UMAP visualization of the pooled real and synthetic cells. miLISI measures the similarity of real and synthetic cells in the UMAP space: the miLISI value ranges between 1 and 2, with 2 indicating a perfect mixing of real and synthetic cells.

**Figure S3:**
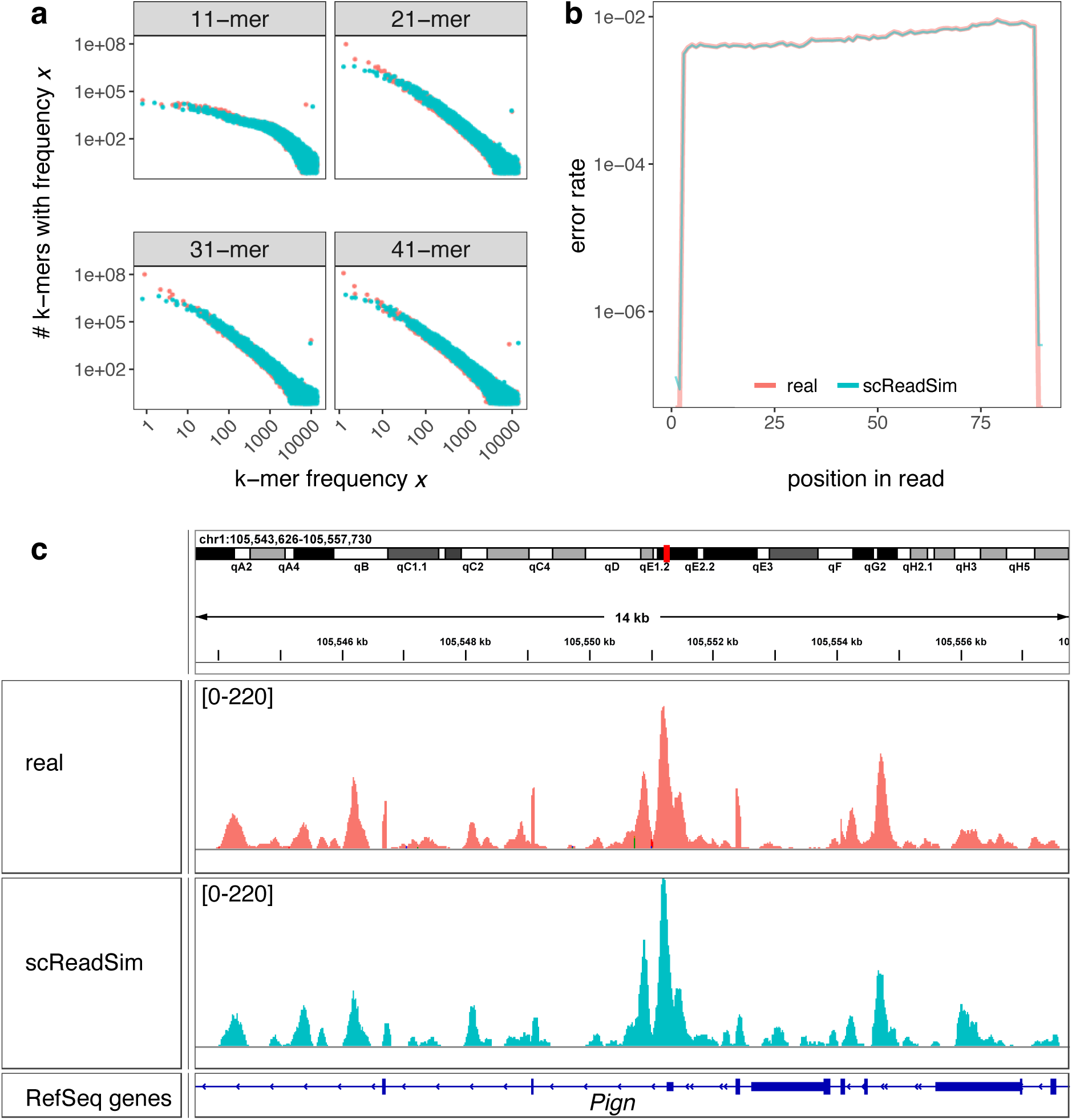
scReadSim’s synthetic data resemble real mouse 10x single-cell Multiome dataset (the RNA-seq modality only) [20] at the read-sequence level. **a**, Comparison of the k-mer spectra between scReadSim’s synthetic data and real data. The x-axis refers to the occurrence frequency of a specific k-mer, and the y-axis represents the number of unique k-mers with this frequency. Both the x-axis and y-axis are in the log_10_ scale. **b**, Comparison of the error rate per base call within a read between scReadSim’s synthetic data and real data. The x-axis represents the positions of bases within a read, and the y-axis refers to the substitution error rate at each position. **c**, Read coverage comparison of the real and synthetic BAM files in the IGV genome browser [25]. The genome browser’s track height is set to 220 for both tracks.

**Figure S4:**
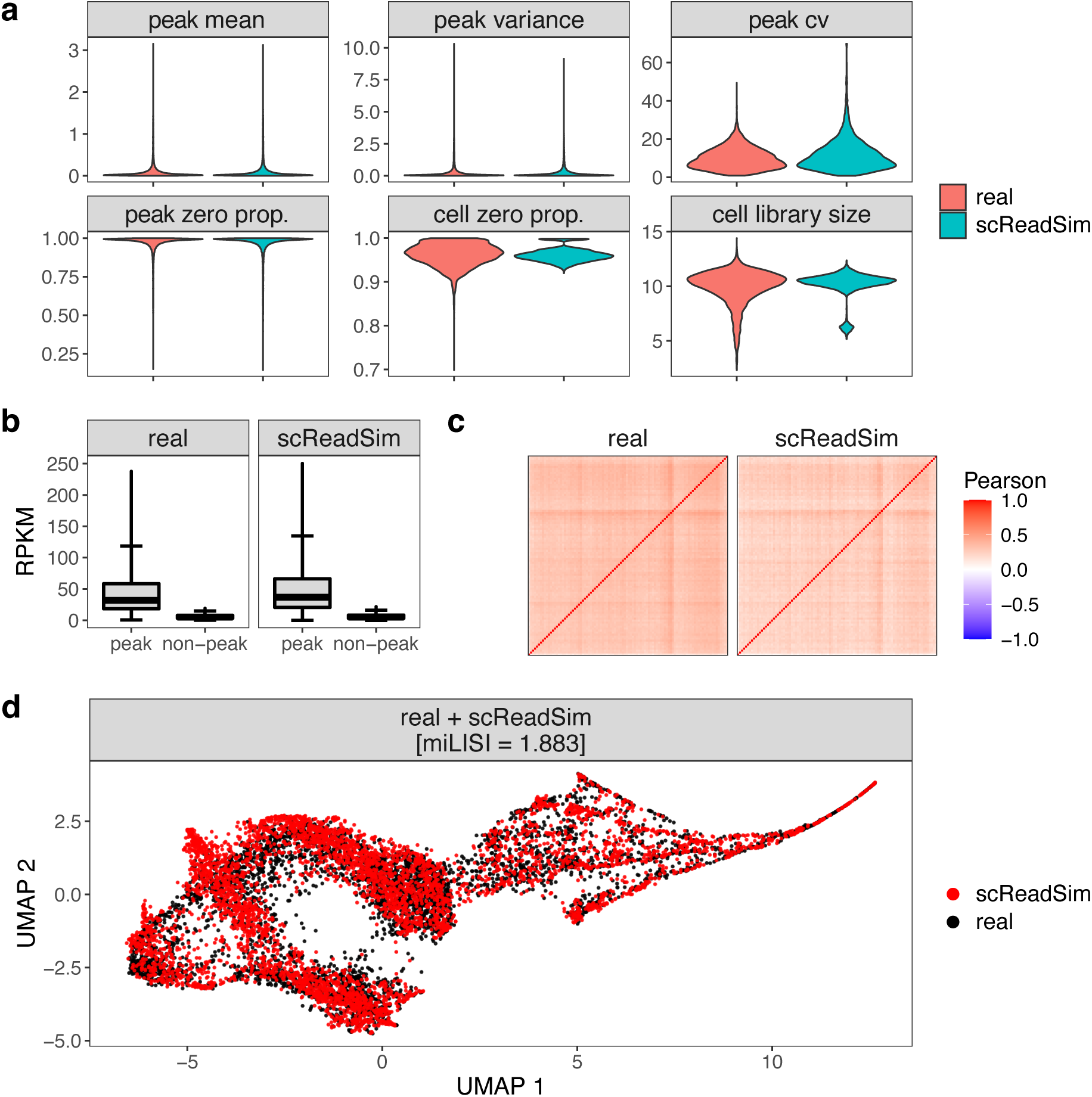
scReadSim’s synthetic data resemble real mouse 10x single-cell Multiome dataset (the ATAC-seq modality only) [20] at the read-count level. **a**, Summary statistics of the count matrix at the peak level (mean, variance, coefficient of variance (cv), zero proportion) and the cell level (zero proportion and cell library size). **b**, Comparison of the RPKM (Reads Per Kilobase Million) value distributions in peak and non-peak regions between scReadSim’s synthetic data and real data. Peaks and non-peaks are obtained using MACS3 from the real BAM file (**Methods**). **c**, Correlations among 100 top peaks in the synthetic and real count matrices. The top-open peaks are defined based on the real count matrix. **d**, UMAP visualization of the pooled real and synthetic cells. miLISI measures the similarity of real and synthetic cells in the UMAP space: the miLISI value ranges between 1 and 2, with 2 indicating a perfect mixing of real and synthetic cells.

**Figure S5:**
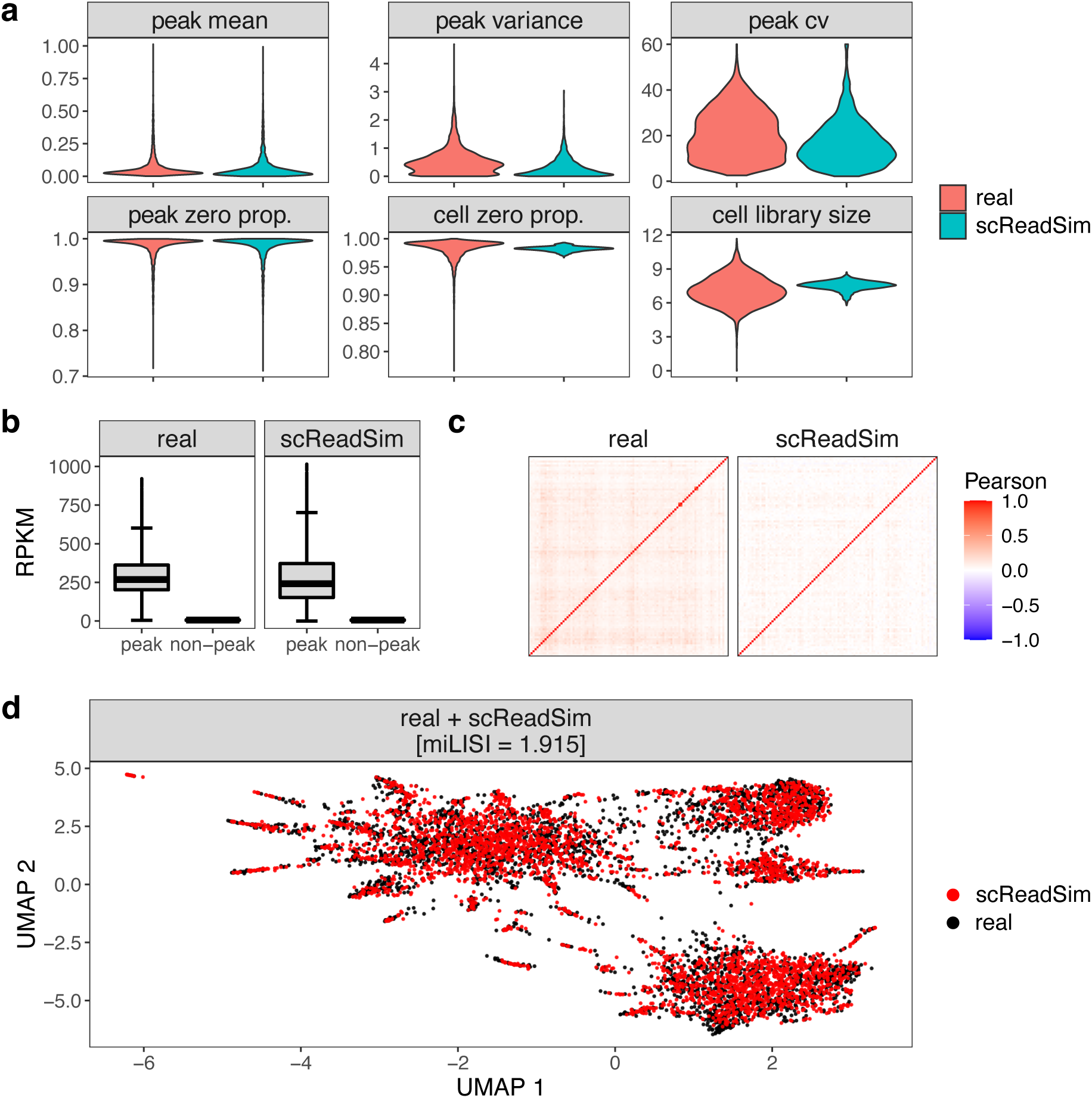
scReadSim’s synthetic data resemble real sci-ATAC-seq data [22]at the read-count level. a, Summary statistics of the count matrix at the peak level (mean, variance, coefficient of variance (cv), zero proportion) and the cell level (zero proportion and cell library size). b, Comparison of the RPKM (Reads Per Kilobase Million) value distributions in peak and non-peak regions between scReadSim’s synthetic data and real data. Peaks and non-peaks are obtained using MACS3 from the real BAM file (Methods). c, Correlations among 100 top peaks in the synthetic and real count matrices. The top-open peaks are defined based on the real count matrix. d, UMAP visualization of the pooled real and synthetic cells. miLISI measures the similarity of real and synthetic cells in the UMAP space: the miLISI value ranges between 1 and 2, with 2 indicating a perfect mixing of real and synthetic cells.

**Figure S6:**
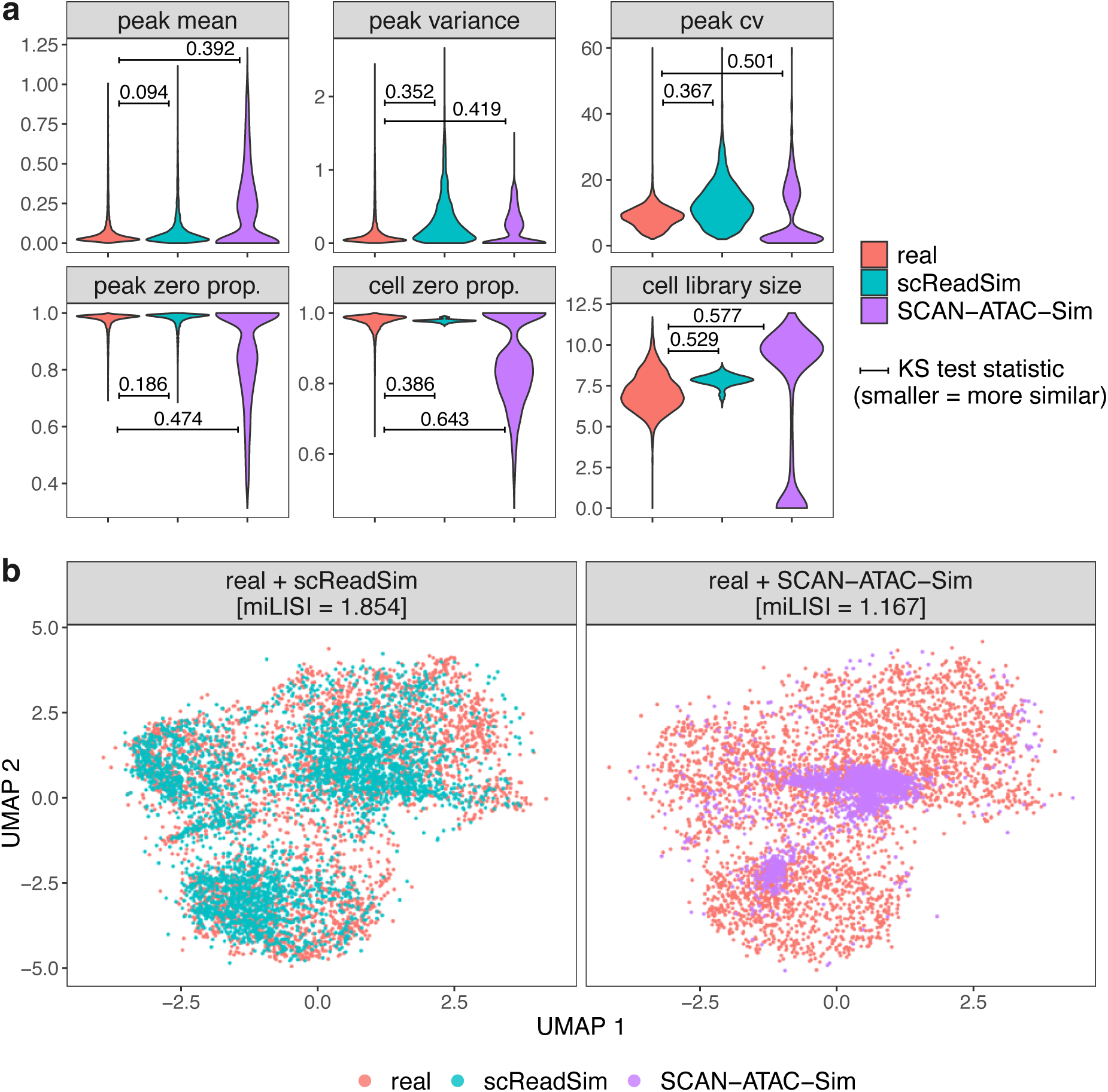
Synthetic cells generated by SCAN-ATAC-Sim do not mimic real cells regarding read counts. **a**, Summary statistics of the count matrix at the peak level (mean, variance, coefficient of variance (cv), zero proportion) and the cell level (zero proportion and cell library size). Kolmogorov–Smirnov tests (KS tests) are performed to compare the empirical distributions of summary statistics between the real and synthetic read count matrices. Smaller KS statistics indicate that the synthetic data mimics real data better. **b**, UMAP visualization of the pooled real and synthetic cells. miLISI measures the similarity of real and synthetic cells in the UMAP space: the miLISI value ranges between 1 and 2, with 2 indicating a perfect mixing of real and synthetic cells.

**Figure S7:**
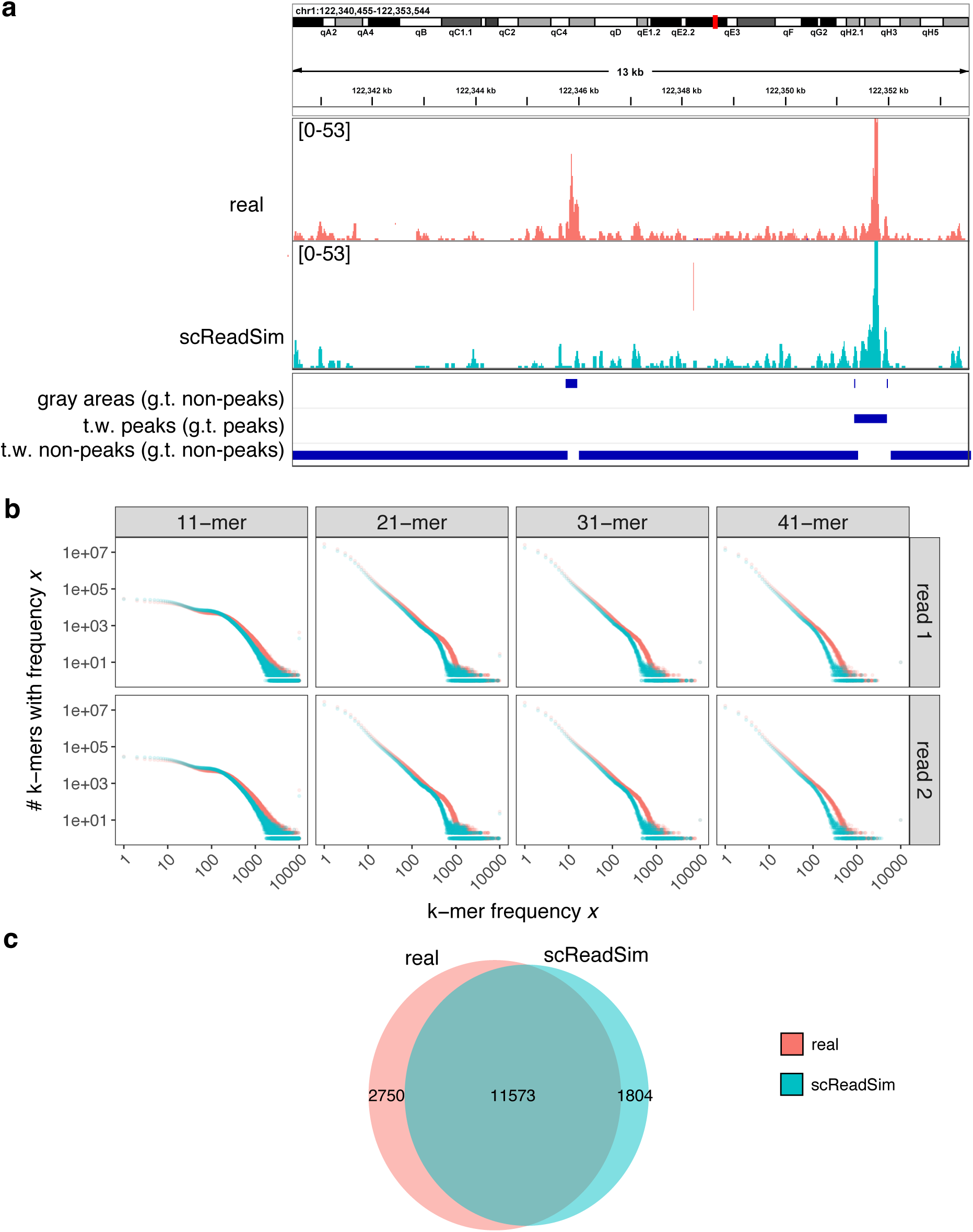
scReadSim’s synthetic data resemble real mouse 10x single-cell Multiome dataset (the ATAC-seq modality only) [20] at the read-sequence level. **a**, Read coverage comparison of the real and synthetic BAM files in the IGV genome browser [25]. The genome browser’s track height is set to 53 for both tracks. Gray areas, trustworthy (t.w.) peaks, and trustworthy non-peaks are user-specified or scReadSim-identified from real data. scReadSim converts gray areas to ground-truth (g.t.) non-peaks, and it maintains trustworthy peaks and non-peaks as ground-truth peaks and non-peaks, respectively, in synthetic data. **b**, Comparison of the k-mer spectra between scReadSim’s synthetic data and real data. The x-axis refers to the occurrence frequency of a specific k-mer, and the y-axis represents the number of unique k-mers with this frequency. Both the x-4ax0is and y-axis are in the log_10_ scale. **c**, A Venn diagram of MACS3’s called peaks from scReadSim’s synthetic data and real data. A peak called from real data (referred to as a “real peak”) is considered to overlap with a peak called from synthetic data (referred to as a “synthetic peak”) if the real peak has at least 101 bp (half of the shortest real peak’s length) overlapped by the synthetic peak.

**Figure S8:**
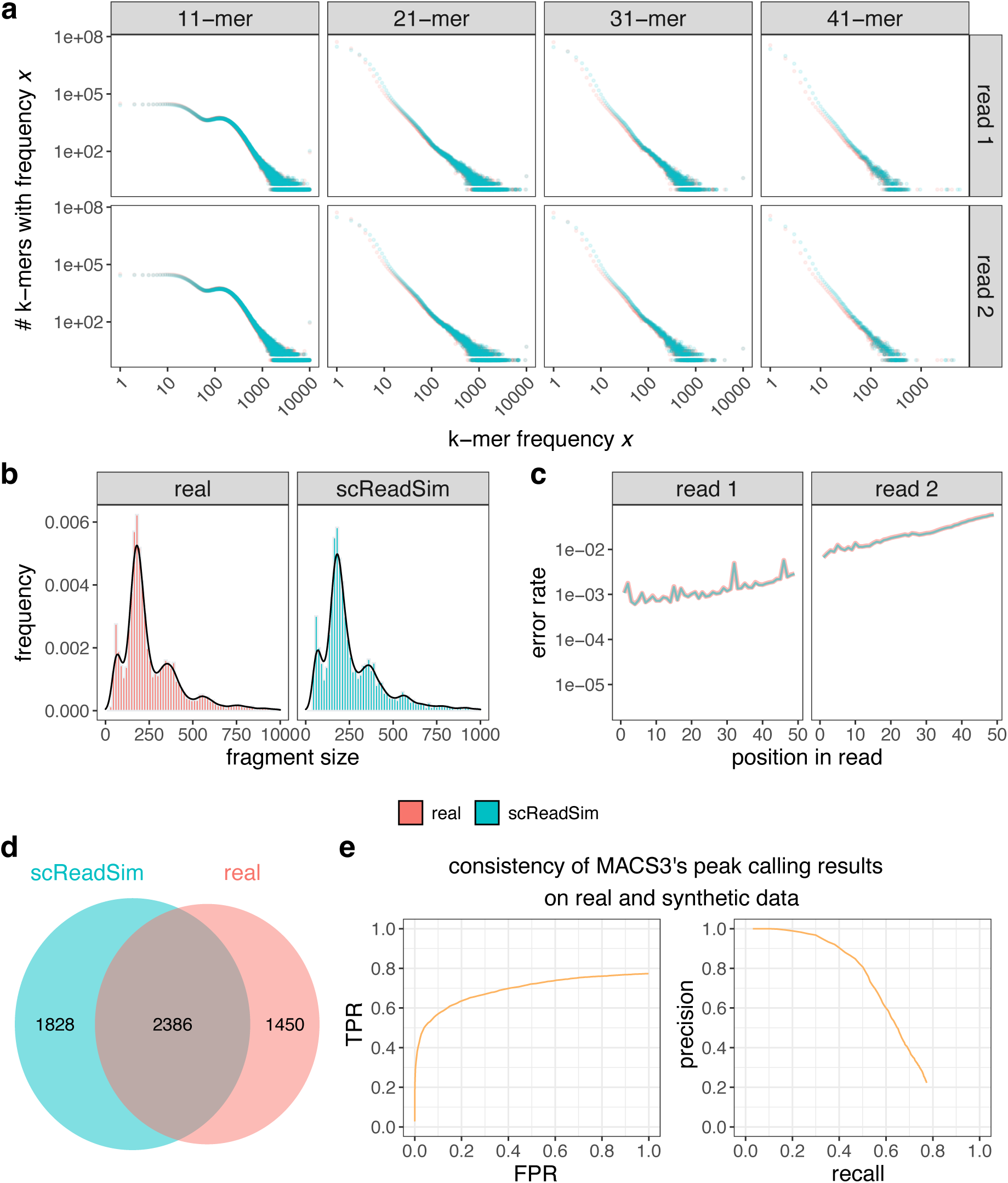
scReadSim’s synthetic data resemble real sci-ATAC-seq data [22] at the read-sequence level. **a**, Comparison of the k-mer spectra between scReadSim’s synthetic data and real data. The x-axis refers to the occurrence frequency of a specific k-mer, and the y-axis represents the number of unique k-mers with this frequency. Both the x-axis and y-axis are in the log_10_ scale. **b**, Comparison of the fragment-size distributions between scReadSim’s synthetic data and real data. **c**, Comparison of the error rate per base call within a read between scReadSim’s synthetic data and real data. The x-axis represents the positions of bases within a read, and the y-axis refers to the substitution error rate at each position. **d**, A Venn diagram of MACS3’s called peaks from scReadSim’s synthetic data and real data. A peak called from real data (referred to as a “real peak”) is considered to overlap with a peak called from synthetic data (referred to as a “synthetic peak”) if the real peak has at least 135 bp (half of the shortest real peak’s length) overlapped by the synthetic peak. **e**, True positive rate (TPR) vs. false positive rate (FPR) (left) and precision vs. recall (right) for evaluating the synthetic peaks in (a) by treating the real peaks in (a) as truths. The TPR, equivalent to the recall, is the proportion of real peaks that have at least 135 bp overlapped by a synthetic peak; the FPR is the proportion of real non-peaks (the regions complementary to the real peaks) that have at least 135 bp overlapped by a synthetic peak; the precision is the proportion of synthetic peaks that overlap at least 135 bp of a real peak.

**Figure S9:**
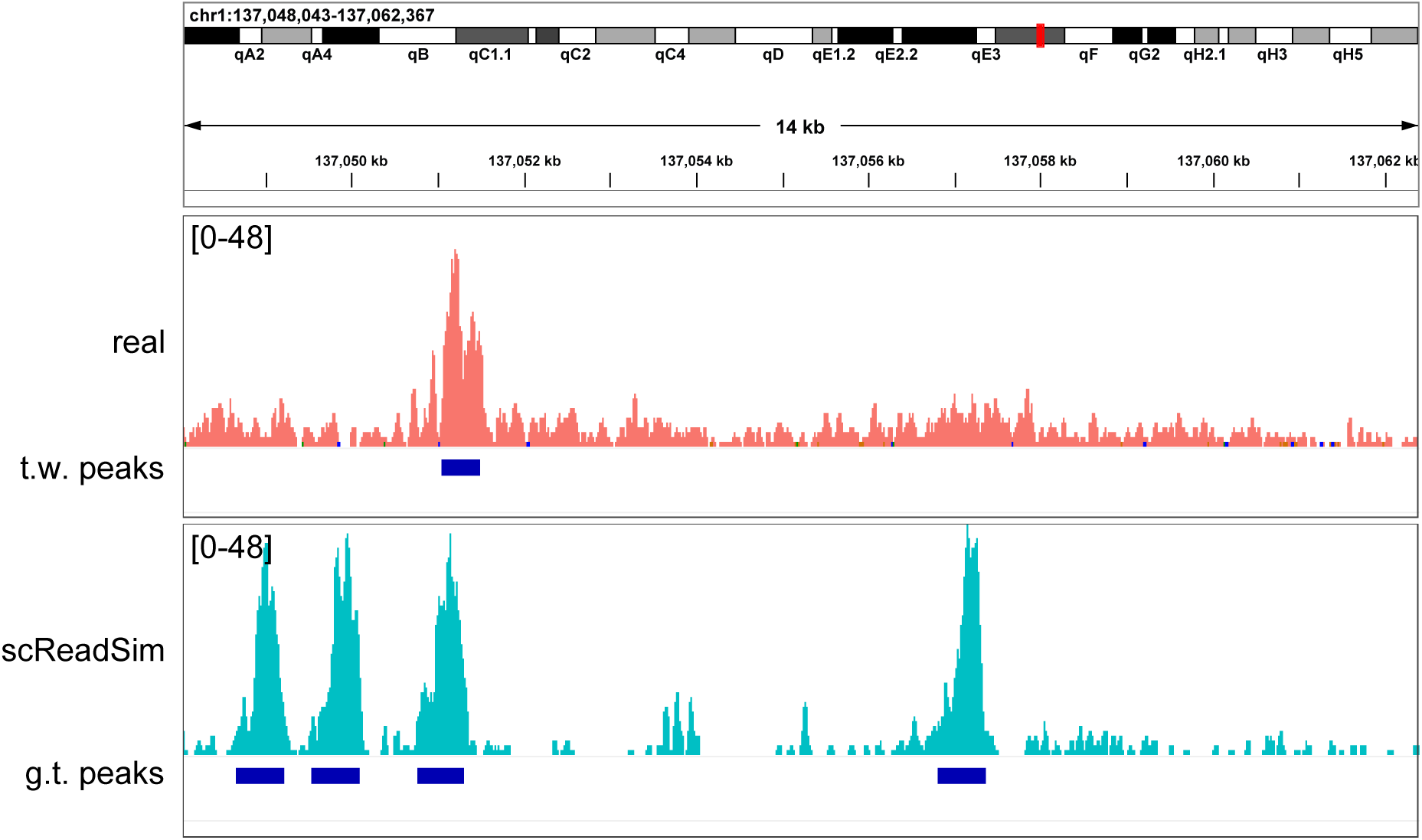
IGV genome browser [25] visualization of scReadSim’s synthetic sci-ATAC-seq reads with user-designed ground-truth (g.t.) peaks (bottom track). From the real sci-ATAC-seq reads (top track) [22], the trustworhy (t.w.) peaks are called by MACS3 and used to train scReadSim. The genome browser’s track height is set to 48 for both tracks.

**Figure S10:**
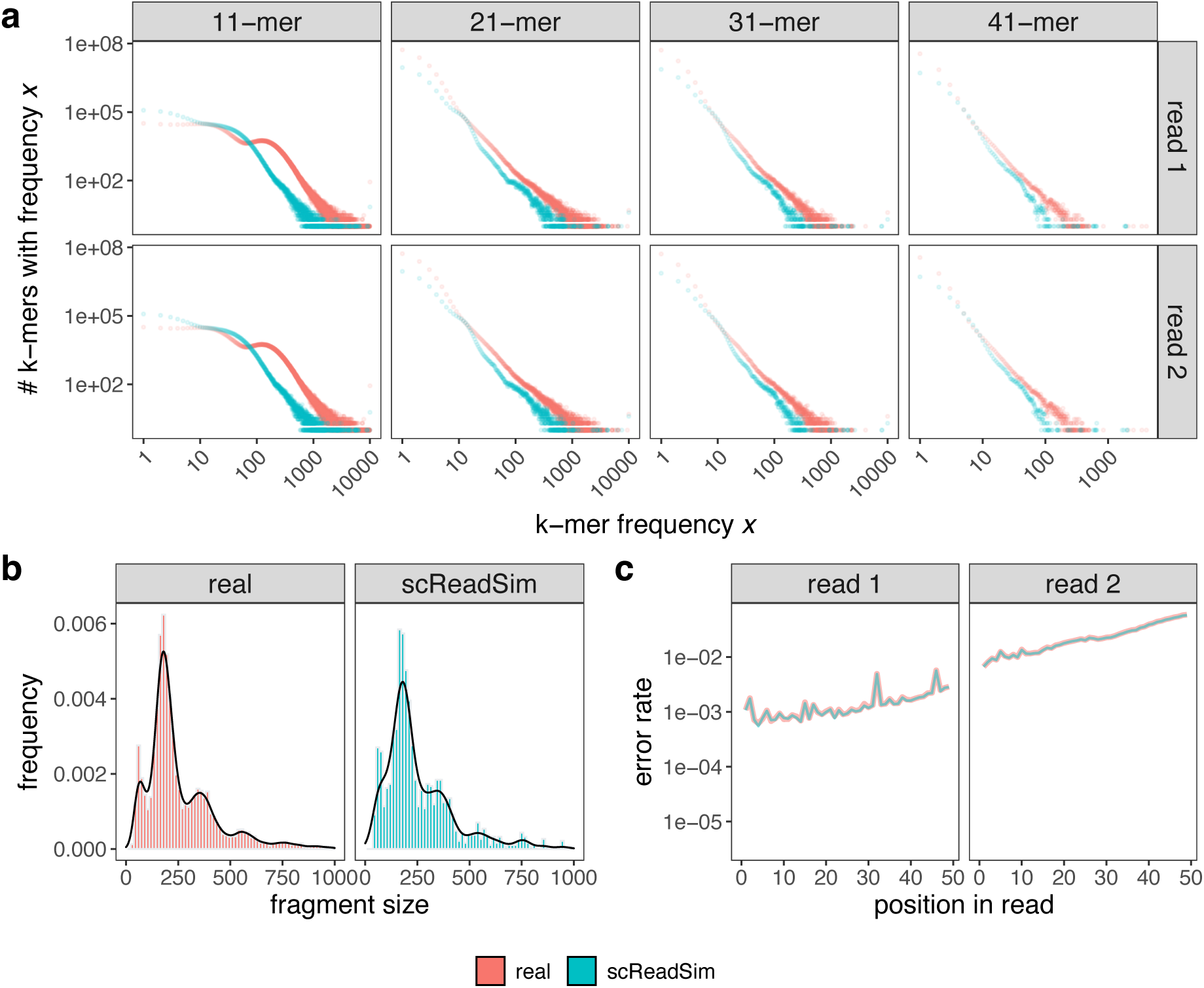
scReadSim’s synthetic data with user-designed open chromatin regions resemble real sci-ATAC-seq data [22] at the read-sequence level. a, Comparison of the k-mer spectra between scReadSim’s synthetic data and real data. The x-axis refers to the occurrence frequency of a specific k-mer, and the y-axis represents the number of unique k-mers with this frequency. Both the x-axis and y-axis are in the log_10_ scale. The real reads and scReadSim’s synthetic reads are expected to have different k-mer spectra because they correspond to different open chromatin regions. b, Comparison of the fragment-size distributions between scReadSim’s synthetic data and real data. c, Comparison of the error rate per base call within a read between scReadSim’s synthetic data and real data. The line plot’s x-axis represents the positions of bases within a read, and the y-axis refers to the substitution error rate corresponding to each position.

**Figure S11:**
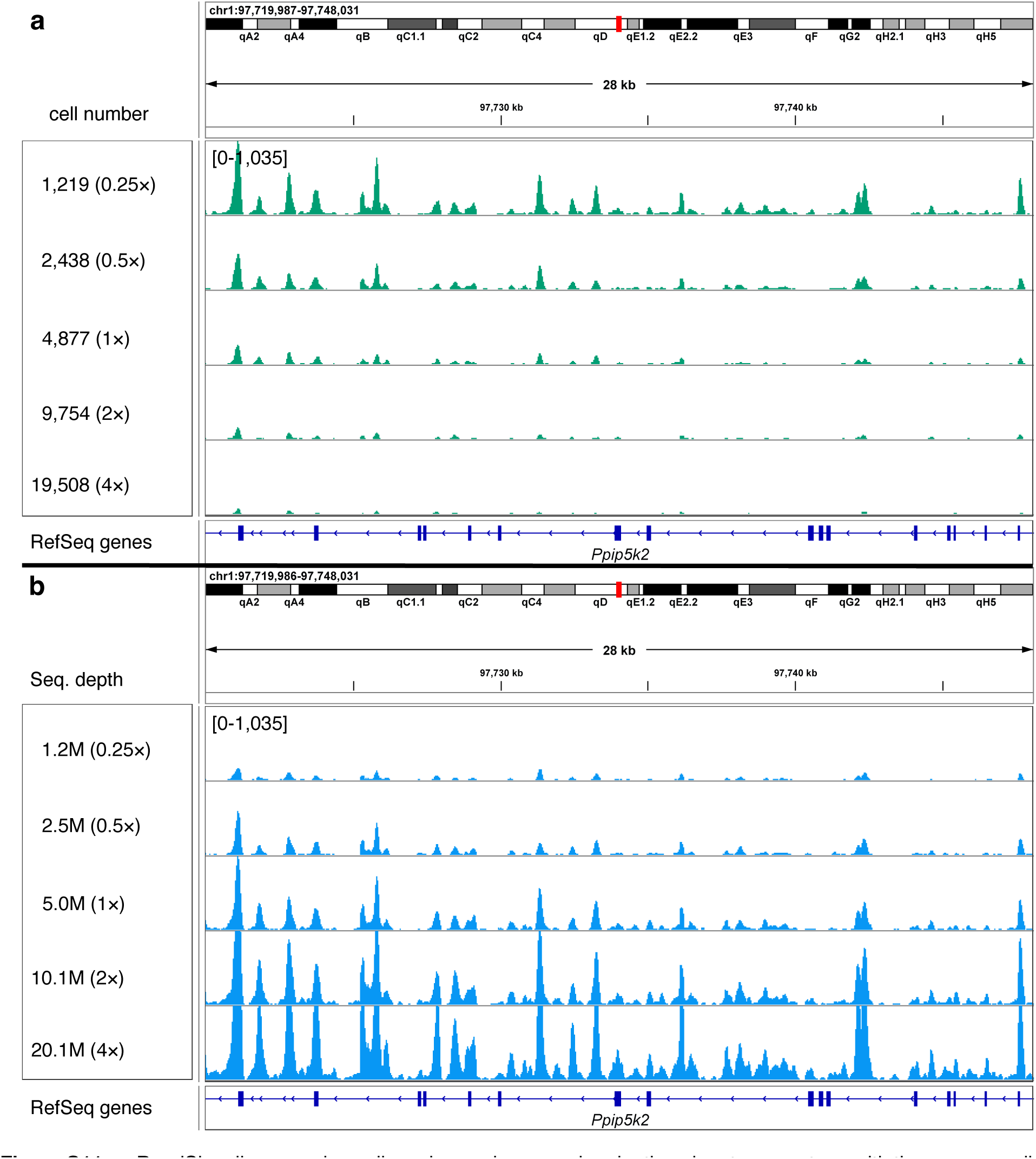
scReadSim allows varying cell number and sequencing depth as input parameters, with the corresponding synthetic data visualized in the IGV genome browser [25]. **a**, Varying numbers of synthetic cells. The track names indicate the multiplication factors: ‘1*×*’ specifies the cell number in the real data; ‘4*×*’ specifies 4 times the real cell number. All five synthetic datasets are downsampled to 1,219 cells for a fair comparison. **b**, Varying sequencing depths in synthetic data. The track names indicate the multiplication factors: ‘1*×*’ specifies the sequencing depth in the real data; ‘4*×*’ specifies 4 times the real sequencing depth. For better visualization, the genome browser’s track height is set to 1,035 for all 10 tracks.

**Figure S12:**
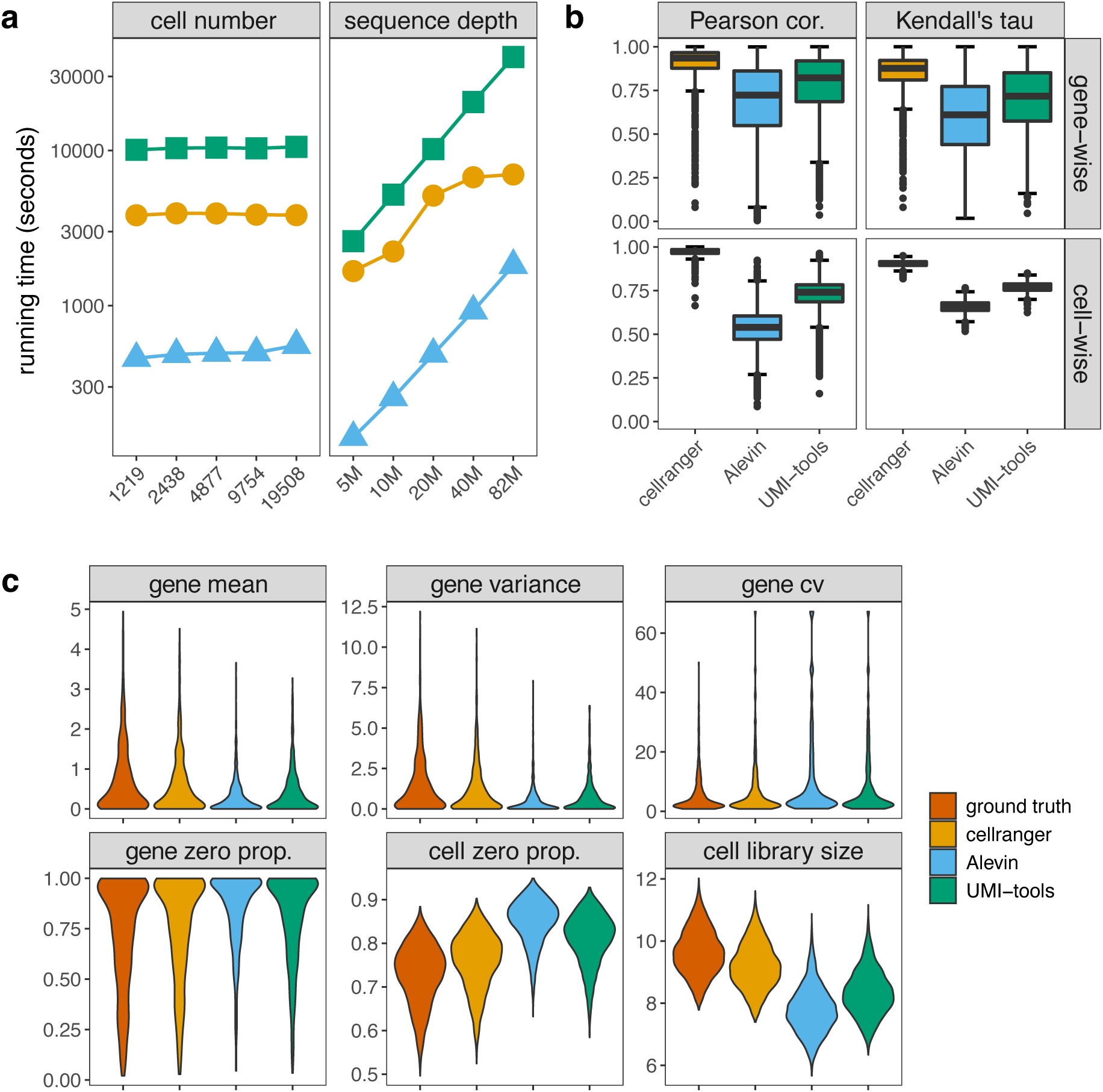
Benchmark of UMI deduplication tools using scReadSim’s synthetic scRNA-seq reads. The ground-truth UMI count matrix is the gene-by-cell UMI count matrix generated by scReadSim. Three deduplication tools are considered: cellranger, Alevin, and UMI-tools. **a**, Time usage of deduplication tools on synthetic datasets with varying cell numbers or sequencing depths. The y-axis indicates the time lapse (in seconds), and the x-axis reflects the number of synthetic cells or the total number of synthetic reads (sequencing depth). **b**, Cell-wise and gene-wise correlations (Pearson correlation and Kendell’s tau) between the ground-truth UMI count matrix and each deduplication tool’s output UMI count matrix. **c**, Summary statistics of the UMI count matrices at the gene level (mean, variance, coefficient of variance (cv), and zero proportion) and the cell level (zero proportion and cell library size).

**Figure S13:**
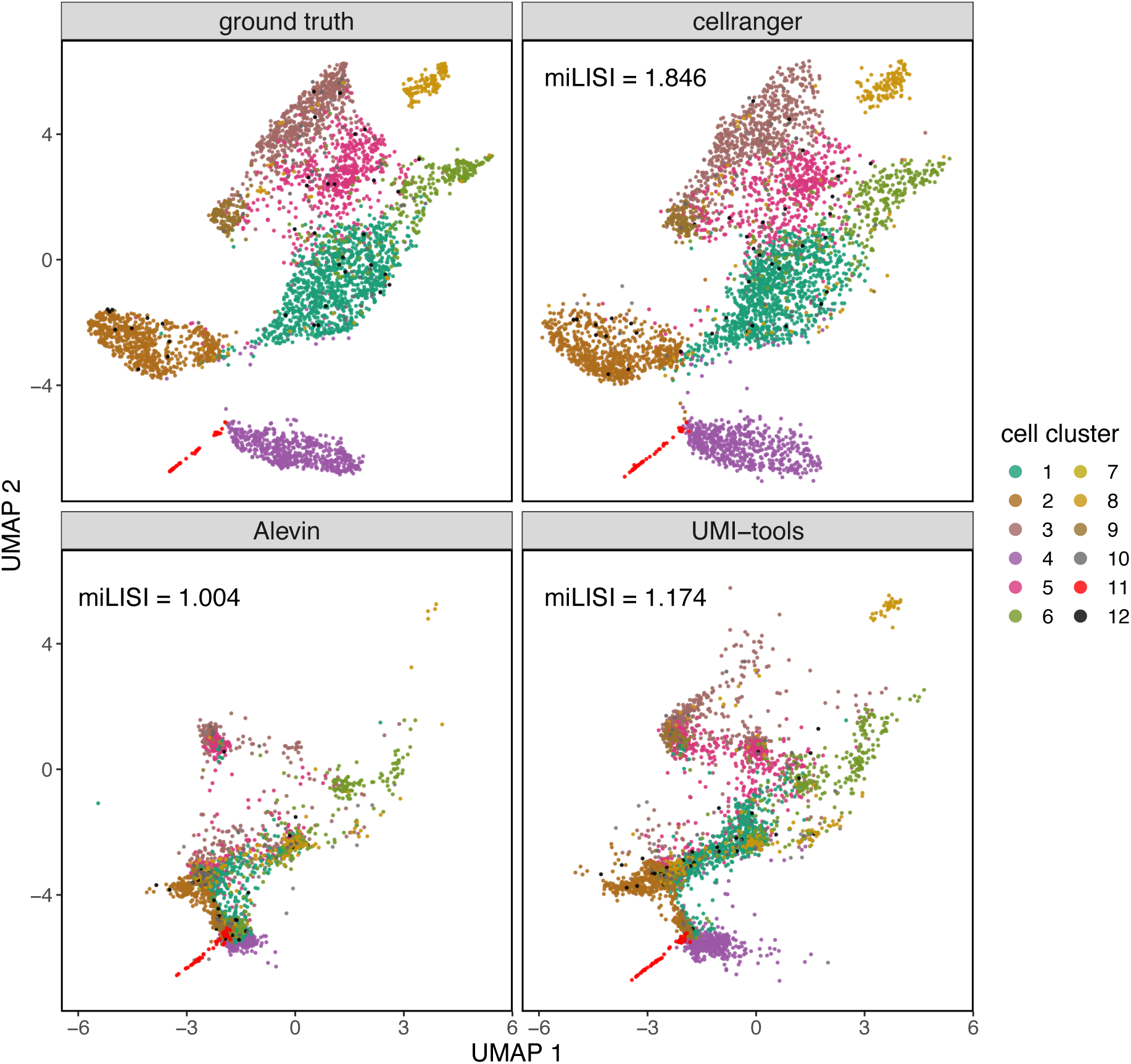
UMAP visualization of the UMI count matrices. Cells are labeled by the cell clusters used as the ground truths in scReadSim. miLISI measures the similarity of real and synthetic cells in the UMAP space: the miLISI value ranges between 1 and 2, with 2 indicating a perfect mixing of real and synthetic cells.

**Figure S14:**
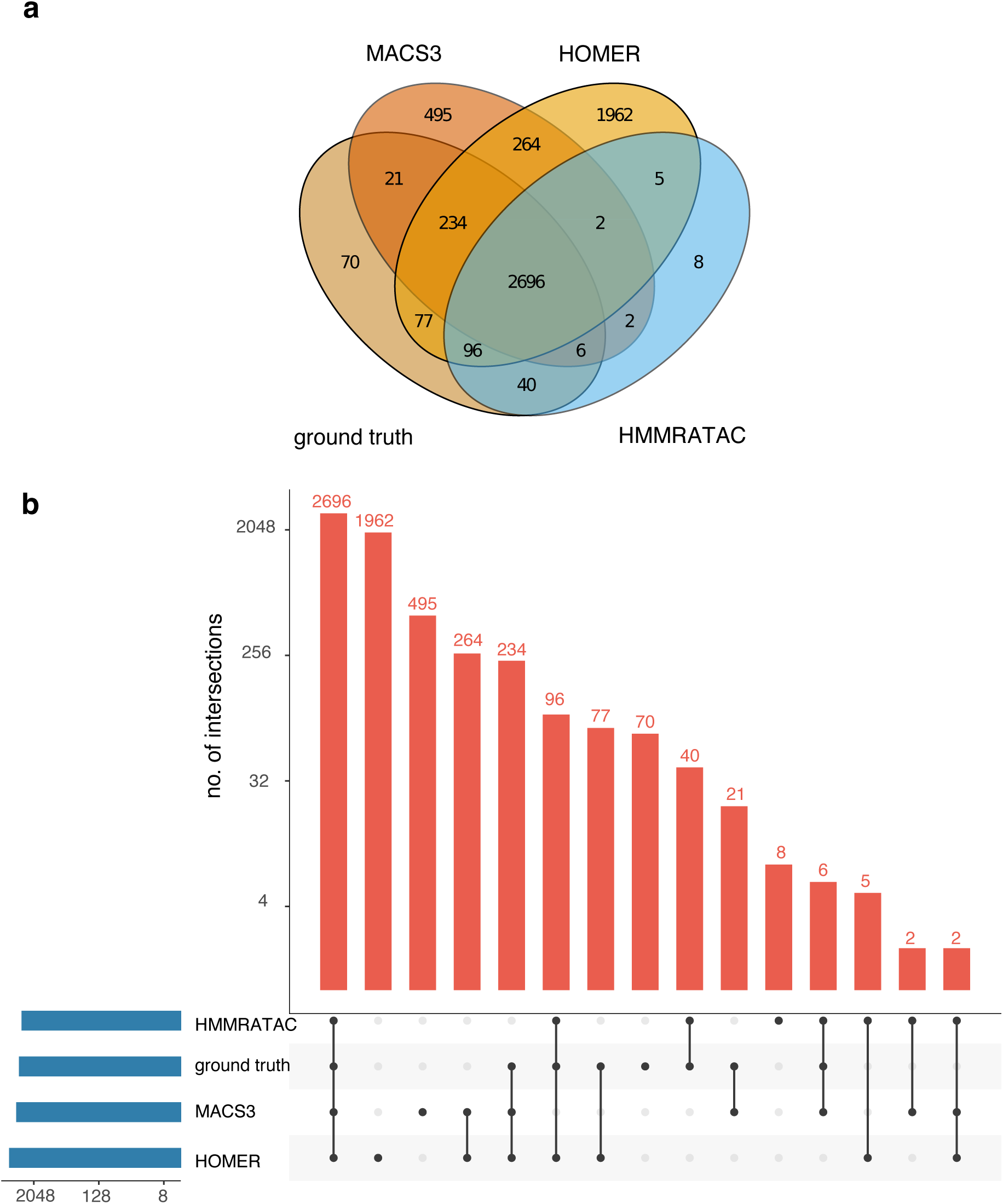
Benchmark of peak-calling tools (MACS3, HOMER, and HMMRATAC) using scReadSim’s synthetic scATAC-seq reads from user-designed ground-truth peaks (i.e., open chromatin regions) (**Methods**). The Venn diagram (**a**) and upset plot (**b**) of the user-designed open chromatin regions and the peaks called by MACS3, HOMER, and HMMRATAC. See Fig. S15 for the comparison results that include the peak-calling tool SEACR, whose called peaks differ drastically from the ground truth and the peaks called by MACS, HOMER, and HMMRATAC.

**Figure S15:**
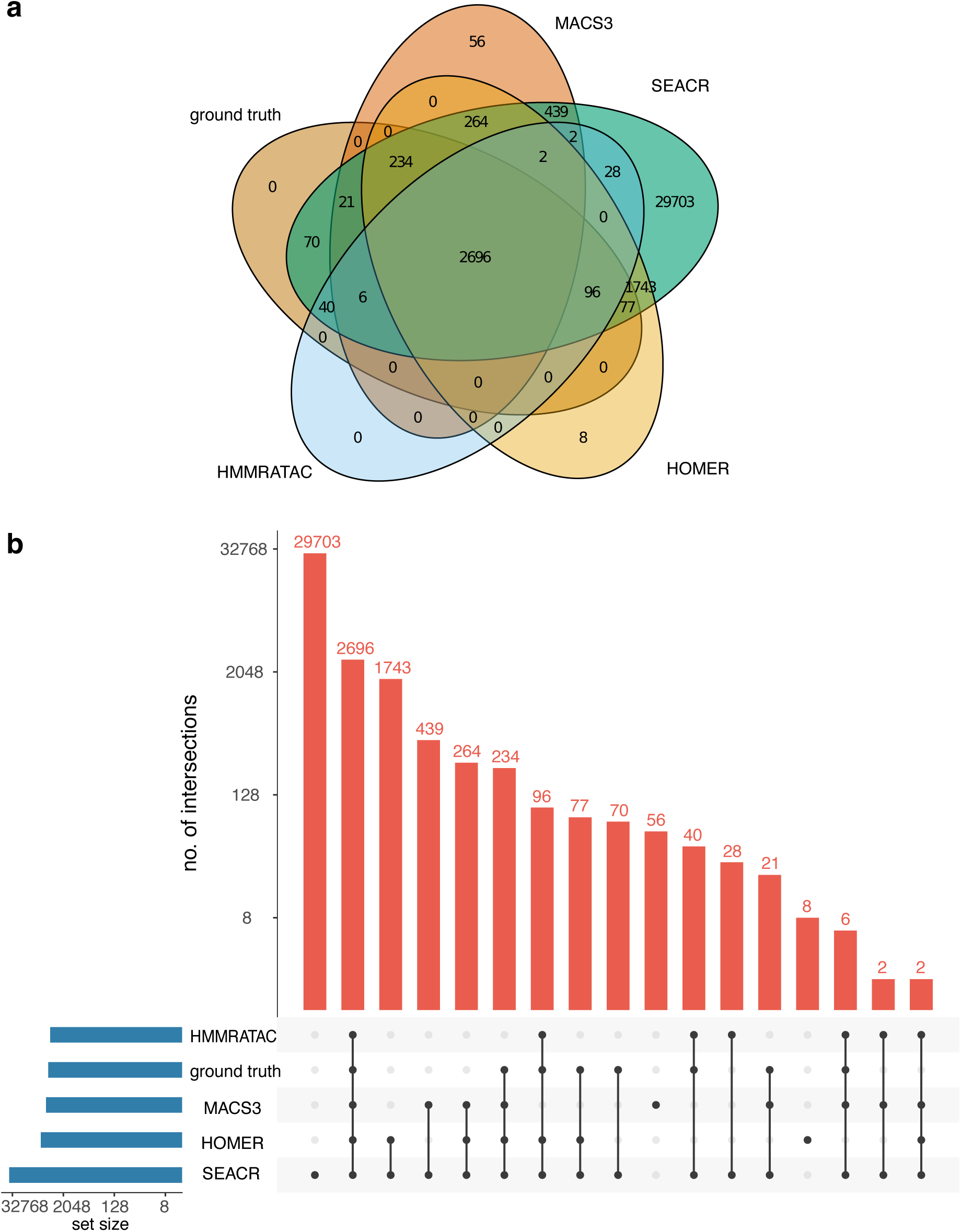
Benchmark of peak-calling tools (MACS3, SEACR, HOMER, and HMMRATAC) using scReadSim’s synthetic scATAC-seq reads from user-designed ground-truth peaks (i.e., open chromatin regions). The Venn diagram (**a**) and upset plot (**b**) of the user-designed open chromatin regions and the peaks called by MACS3, SEACR, HOMER, and HMMRATAC.

**Figure S16:**
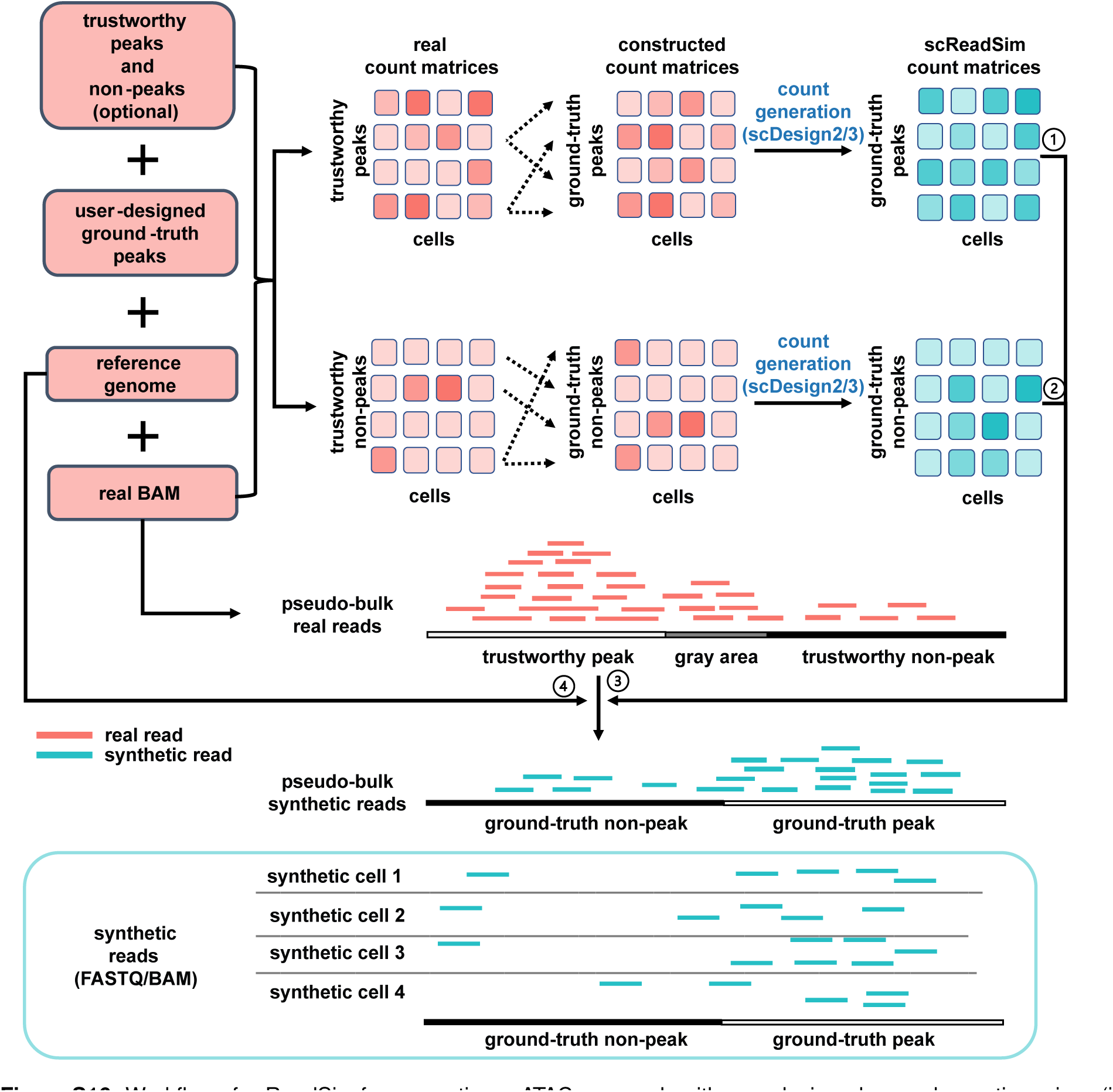
Workflow of scReadSim for generating scATAC-seq reads with user-designed open chromatin regions (i.e., ground-truth peaks). The input includes a BAM file, the corresponding reference genome, trustworthy peak and non-peak lists, and a ground-truth peak list that the synthetic scATAC-seq reads will be generated accordingly. Specifically, if users do not specify the trustworthy peaks and non-peaks, scReadSim by default uses MACS3 [13] with stringent criteria to call trustworthy peaks (q-value *≤* 0.01) and non-peaks (q-value *>* 0.1) from the input BAM file. Given ground-truth peaks, scReadSim defines the inter-peaks as the ground-truth non-peaks. Based on trustworthy peaks and non-peaks, scReadSim summarizes scATAC-seq reads in the input BAM file into a trustworthy-peak-by-cell count matrix and a trustworthy-non-peak-by-cell count matrix. Next, scReadSim defines a mapping between trustworthy and ground-truth peaks and constructs a ground-truth-peak-by-cell count matrix by extracting the corresponding trustworthy-peak-by-cell count matrix’s entries. Similarly, scReadSim constructs a ground-truth-non-peak-by-cell count matrix from the non-trustworthy-peak-by-cell count matrix. Further, scReadSim trains the count simulator scDesign2 [6] (if the cells belong to distinct clusters; otherwise, scDesign3 [21] can be used if the cells follow continuous trajectories) on the ground-truth-peak- and ground-truth-non-peak-by-cell count matrices to generate the corresponding synthetic count matrices for the ground-truth peaks and non-peaks. Last, scReadSim generates synthetic reads based on the synthetic count matrices (*0*1 and *0*2), the input BAM file (*0*3), and the reference genome (*0*4). The synthetic reads are outputted in FASTQ and BAM format.

**Figure S17:**
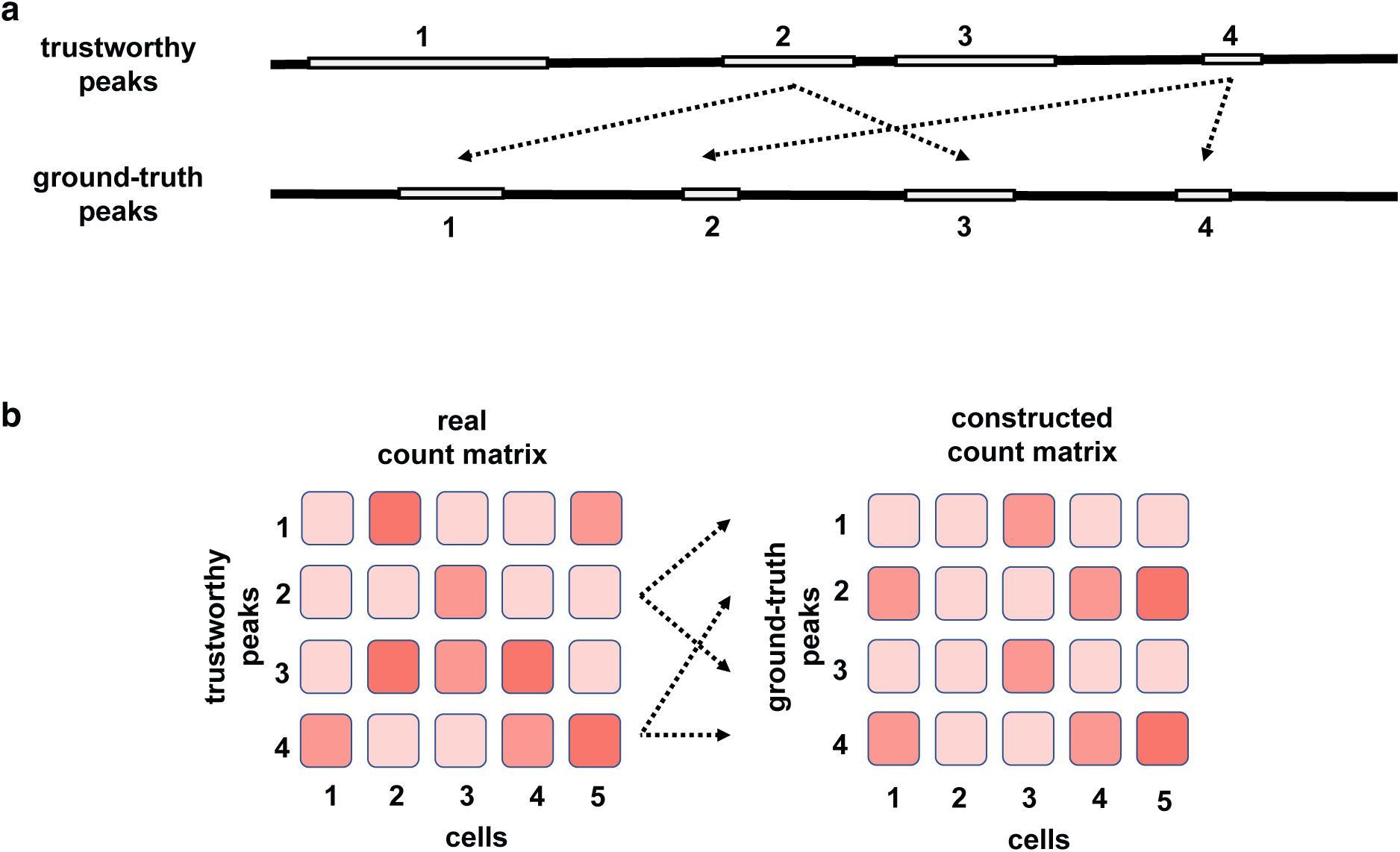
The mechanism of scReadSim’s data generation with user-designed open chromatin regions (i.e., ground-truth peaks). a, The mapping between trustworthy peaks and ground-truth peaks. For every ground-truth peak, scReadSim finds its most similar trustworthy peak (in the same chromosome) in terms of length. b, The constructed count matrix for ground-truth peaks. For every ground-truth peak, scReadSim adds the corresponding trustworthy peak’s row in the trustworthy peak-by-cell count matrix to the ground-truth peak-by-cell count matrix.

## Supplementary Tables

**Table S1:** Comparison of scReadSim, minnow, and SCAN-ATAC-Sim

| **Tools** | **scReadSim** | **minnow** | **SCAN-ATAC-Sim** |
| --- | --- | --- | --- |
| **Property** |  |  |  |
| Modality | scRNA-seq<br>scATAC-seq | scRNA-seq | scATAC-seq |
| Input | scRNA-seq reads<br>or scATAC-seq reads | UMI count matrix | ATAC-seq reads<br>peak lists |
| Output | FASTQ<br>or BAM | FASTQ | BED |
| Ground truth | UMI counts<br>open chromatin regions | UMI counts | N/A |
| Read-sequence preservation<sup>1</sup> | ✓ | ✗<sup>2</sup> | ✗ |
| Read-count preservation<sup>3</sup> | ✓ | ✓ | ✗ |
| Read-coverage preservation<sup>4</sup> | ✓ | ✗ | ✗<sup>5</sup> |
| Whole-genome read simulation | ✓ | ✗ | ✓ |

<sup>1</sup>: “Read-sequence preservation” means whether the simulator preserves the characteristics of real read sequences, such as k-mer spectra and sequencing errors.
<sup>2</sup>: Since minnow does not input scRNA-seq reads, it uses its internal, pre-trained model to generate sequencing errors.
<sup>3</sup>: “Read-count preservation” means whether the simulator generates synthetic cells that mimic real cells regarding gene or peak counts.
<sup>4</sup>: “Read-coverage preservation” means whether the simulator’s synthetic reads have a coverage profile mimicking that of real reads at the pseudo-bulk level.
<sup>5</sup>: SCAN-ATAC-Sim only inputs pseudo-bulk ATAC-seq reads and thus does not preserve read coverage at the single-cell level.

## Methods

### The scReadSim method

The input of scReadSim is a BAM file storing the aligned sequencing reads from scRNA-seq or scATAC-seq. As an overview, scReadSim consists of three major steps: genome segmentation, synthetic count generation, and synthetic read generation.

The first step segments the genome so that each genomic region (segment) is defined as a feature. Specifically, for scRNA-seq, scReadSim requires users to input a gene annotation file to segregate the genome into genes and inter-genes; for scATAC-seq, scReadSim requires trustworthy peaks and non-peaks (if not user-specified (optional), scReadSim by default uses MACS3 with stringent criteria to call trustworthy peaks (q-value *≤* 0.01) and non-peaks (q-value *>* 0.1) from the input BAM file) and defines the complementary genomic regions as gray areas.

Next, scReadSim summarizes scRNA-seq reads in the input BAM file into a gene-by-cell UMI count matrix and an inter-gene-by-cell UMI count matrix, or scReadSim summarizes scATAC-seq reads in the input BAM file into a peak-by-cell count matrix and a non-peak-by-cell count matrix. We use a general term “feature-by-cell count matrix” to refer to each count matrix in the following text.

In the second step—synthetic count generation, scReadSim trains the count simulator scDesign2 [6] (if the cells belong to distinct clusters; otherwise, scDesign3 [21] can be used if the cells follow continuous trajectories) on the feature-by-cell count matrices to generate the corresponding synthetic count matrices. For scATAC-seq, scReadSim converts gray areas into non-peaks before generating the corresponding synthetic count matrix. In summary, scReadSim generates two synthetic count matrices—gene-by-cell and inter-gene-by-cell matrices—for scRNA-seq; scReadSim generates three synthetic count matrices—peak-by-cell, non-peak-by-cell, and non-peak-by-cell (converted from gray areas) matrices—for scATAC-seq.

The third step generates synthetic reads based on the synthetic count matrices, the input BAM file, and the reference genome. The synthetic reads are outputted in FASTQ and BAM formats. Fig. 1a–b illustrate the scReadSim workflows for scRNA-seq and scATAC-seq, respectively.

### scReadSim for scRNA-seq

The procedure below is for simulating data from UMI-based scRNA-seq technologies such as 10x Chromium [26] and Drop-seq [27]. For non-UMI-based scRNA-seq technologies such as Smart-seq2 [28], the simulation procedure follows the next subsection “scReadSim for scATAC-seq with user-designed open chromatin regions” with user-designed open chromatin regions replaced by genes.

1. Pre-processing real scRNA-seq data to obtain UMI-count matrices

To pre-process real scRNA-seq data for training, scReadSim requires a BAM file (containing scRNA-seq reads in cells) and a gene annotation file (in GTF format). Based on the gene coordinates in the annotation file, scReadSim segregates the reference genome into two sets of features: genes and inter-genes. Then scReadSim counts the number of UMIs whose reads overlap with each gene in every cell to construct a gene-by-cell UMI-count matrix, and similarly, an inter-gene-by-cell UMI-count matrix. The two matrices contain the same cells in the same column order.

2. Synthetic UMI-count matrix generation

To generate the gene- and inter-gene-by-cell UMI-count matrices for synthetic cells, scReadSim trains the count simulator scDesign2 [6] on the pre-processed gene- and inter-gene-by-cell UMI-count matrices from real data, respectively. Note that scReadSim chooses scDesign2 because scDesign2 can generate realistic synthetic counts by capturing gene-gene correlations [9]; however, scReadSim is flexible to accommodate any scRNA-seq count simulator.

Specifically, scDesign2involves two steps: (1) model fitting on the real data count matrix and (2) generation of a synthetic count matrix by sampling synthetic cells from the fitted model. Hence, scReadSim uses scDesign2 to generate synthetic gene-by-cell and inter-gene-by-cell count matrices from the real gene-by-cell and inter-gene-by-cell count matrices, respectively. Since scDesign2 requires cells to be in discrete cell types, scReadSim addresses this requirement by pre-clustering the cells using the gene-by-cell count matrix and Louvain clustering (in the Seurat R package [29]) if cells do not have pre-defined cell type labels. Note that scReadSim can also use the updated count simulator scDesign3 [21] to generate synthetic count matrices if cells are from continuous trajectories instead of discrete cell types.

In summary, scReadSim generates the synthetic gene-by-cell and inter-gene-by-cell UMI-count matrices in this step.

3. Synthetic read generation

To generate the scRNA-seq reads for synthetic cells, scReadSim uses:

- the synthetic gene-by-cell and inter-gene-by-cell UMI-count matrices,
- the coordinates of the genes and inter-genes,
- the input BAM file,
- the reference genome,
- the user-specified read length (optional; default 90 nt).

In UMI-based scRNA-seq data, every paired-end read has only one end (referred to as read 2) containing RNA sequence information, while the other end (referred to as read 1) contains cell barcode and UMI information [17, 30]. The generation of these two ends is described below (Fig. 1a).

(a) Generation of read 2 (RNA sequence)

For simplicity, we use a “read” to refer to a read 2. For each gene or inter-gene *i* in each synthetic cell, scReadSim (1) extracts the corresponding UMI count *c_u_* (i.e., the number of synthetic UMIs) from the synthetic feature-by-cell UMI-count matrices;

(2) summarizes the read count of each UMI from the real reads that overlap with this feature and obtains the empirical distribution *F_i_* of the read count per UMI; (3) samples the number of synthetic reads *c* for each synthetic UMI according to the empirical distribution *F*^^^*_i_*; and (4) for each synthetic UMI with read count *c*,

i. samples with replacement a real UMI from the real UMIs with read counts equal to *c*;
ii. samples (with replacement) *c* real reads from those that are associated with the sampled real UMI;
iii. generates *c* synthetic reads from the *c* real reads, the read length, and the reference genome;
iv. assigns the *c* synthetic reads to the synthetic UMI.

Specifically, in (4), scReadSim converts the *c* real reads’ 5’ positions in the reference genome into the *c* synthetic reads’ 5’ positions (after adding random shifts, detailed in the next paragraph); then scReadSim finds the *c* synthetic reads’ 3’ positions in the reference genome based on the read length (e.g., if a synthetic read has the 5’ position *x* and the read length *l*, then its 3’ position is *x*+*l−*1). Next, given every synthetic read’s 5’ and 3’ positions, and strand information (preserved from real reads), scReadSim extracts the read sequence from the reference genome sequence.

Regarding the random shifts mentioned in the above paragraph, scReadSim adds a random shift (sampled uniformly from *−*4 to 4 nt) to each real read’s 5’ position to obtain a synthetic read’s 5’ position. The purpose of doing so is to resolve duplicate 5’ positions due to sampling with replacement. To make the synthetic reads more realistic, scReadSim also allows the introduction of substitution errors, as described in subsection “Substitution errors introduced into synthetic reads.”

(b) Generation of read 1 (cell barcode and UMI)

Read 1 is a concatenated sequence string of a UMI and a cell barcode. Accordingly, scReadSim concatenates a randomly generated cell barcode and a UMI to form a synthetic read 1, which will then be paired with a synthetic read 2 generated above. Below we describe the generation of a cell barcode and a UMI for each synthetic read 1, whose length is specified as 26 nt (including a 16-nt cell barcode and a 10-nt UMI).

Specifically, for each synthetic cell, scReadSim generates a 16-nt cell barcode by randomly sampling A, C, G, and T with replacement for 16 times and assigns this cell barcode to all synthetic reads belonging to the synthetic cell. To generate a UMI for each gene or inter-gene in each synthetic cell, scReadSim (1) extracts the corresponding UMI count *c_u_* (i.e., the number of UMIs) from the synthetic feature-by-cell UMI-count matrices; (2) generates *c_u_* 10-nt UMIs, with each UMI created by randomly sampling A, C, G, and T with replacement for 10 times; and (3) assigns the *c_u_* UMIs to the synthetic read 2 based on the read-UMI relationship generated above. Note that the lengths of cell barcodes and UMIs can be user-specified (default lengths are 16 and 10 nt, respectively).

In summary, scReadSim outputs paired-end synthetic reads in two FASTQ files for read 1 and read 2 separately. Regarding the Phred quality scores in the FASTQ files, the read 2 FASTQ file uses the quality score 37 for reference sequences and the quality score 24 for erroneous substitutions, while the read 1 FASTQ file uses the quality score 37 for all sequences. Optionally, scReadSim can output a BAM file, which contains the mapped reads from the two FASTQ files to the reference genome.

### scReadSim for scATAC-seq

1. Pre-processing

To pre-process real scATAC-seq data for training, scReadSim requires a BAM file (containing scATAC-seq reads in cells) and users’ trustworthy peaks and non-peaks in the input BAM file. Alternatively, if users do not specify trustworthy peaks and non-peaks, scReadSim first deploys the peak-calling tool MACS3 [13] to identify the trustworthy peaks using a stringent rule (by setting a q-value threshold 0.01) and the trustworthy non-peaks using a loose rule (by setting a q-value threshold 0.1; the non-peaks are complementary to the peaks called under this threshold).

Based on the trustworthy peaks and non-peaks, scReadSim defines “gray areas” as the genomic regions complementary to the peaks and non-peaks, and such gray areas represent the chromatin regions that cannot be confidently classified as peaks or non-peaks. In summary, scReadSim segments the reference genome into three sets of features: peaks, non-peaks, and gray areas (Fig. 1b). Then scReadSim counts the number of reads overlapping each peak in every cell to construct a peak-by-cell count matrix. scReadSim also generates a non-peak-by-cell count matrix similarly. The two matrices contain the same cells in the same column order.

2. Synthetic count matrix generation

In scATAC-seq read generation, scReadSim provides ground-truth peaks and non-peaks for benchmarking purposes by learning from user-specified trustworthy peaks and non-peaks or calling trustworthy peaks and non-peaks from real data. To generate the peak-by-cell and non-peak-by-cell count matrices for synthetic cells, scReadSim trains the count simulator scDesign2 on the pre-processed peak-by-cell and non-peak-by-cell count matrices from real data. Next, scReadSim converts the gray areas into non-peaks (so that the peaks can be regarded as “ground-truth peaks”) and constructs a synthetic count matrix based on the gray areas’ lengths, trustworthy non-peaks’ lengths, and the previous synthetic non-peak-by-cell count matrix.

Specifically, scReadSim generates gray areas’ synthetic read counts based on the trustworthy non-peaks’ synthetic read counts so that the gray areas’ synthetic read coverage mimics the non-peaks’ synthetic read coverage. For each gray area,

(a) scReadSim first finds the closest non-peak on the 5’ of the gray area and extracts the non-peak’s synthetic count vector (the non-peak’s corresponding row) from the synthetic non-peak-by-cell count matrix;

(b) scReadSim then randomly masks (sets to 0) the count entries of the non-peak’s synthetic count vector with a probability equal to 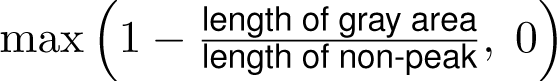 and assigns the masked count vector as the gray area’s synthetic count vector.

This random masking essentially scales the non-peak’s synthetic read coverage according to the ratio of the gray area’s length and the non-peak’s length so that the gray area’s synthetic read coverage mimics that of its nearby non-peak.

3. Synthetic read generation

To generate the scATAC-seq reads for synthetic cells, scReadSim uses

- the synthetic peak-by-cell, non-peak-by-cell count matrices,
- the coordinates of the peaks and non-peaks,
- the input BAM file,
- the reference genome,
- the user-specified read length (optional; default 50 nt).

As scATAC-seq usually generates paired-end reads, scReadSim accordingly generates synthetic paired-end reads. For clarity, we use a “read” to denote one end in a paired-end read. For each peak or non-peak in each synthetic cell, scReadSim (1) extracts the corresponding count *c* (i.e., the number of synthetic reads) from the synthetic feature-by-cell (feature refers to peak or non-peak) count matrices; (2) samples (with replacement) *Ic/*2*1* (rounding *c/*2 to the least integer greater than or equal to *c/*2) real reads that overlap this feature in the BAM file; (3) finds the mates of (i.e., real reads in pairs with) the *Ic/*2*1* sampled real reads from the BAM file; and (4) generates *Ic/*2*1* synthetic read pairs from the *Ic/*2*1* real read pairs, the read length, and the reference genome.

Specifically, in (4), scReadSim follows a similar procedure to the one in [subsection “scReadSim for scRNA-seq” *→* “3. Synthetic read generation” *→* “(a) Generation of read 2 (RNA sequence)” *→* “step (4) iii. generates *c* synthetic reads from the *c* real reads, the read length, and the reference genome”]. The only difference is the accommodation of paired-end reads.

In the random shifting step to avoid duplicate reads, scReadSim adds the same random shift to a pair of reads’ 5’ positions to keep the fragment size unchanged after the shift.

Typically, a scATAC-seq BAM file stores cell barcodes as additional information. To generate a cell barcode (e.g., 16-nt long) for each synthetic cell, scReadSim randomly samples A, C, G, and T with replacement 16 times, and assigns this cell barcode to all synthetic reads belonging to the synthetic cell.

scReadSim for scATAC-seq with user-designed open chromatin regions

1. Pre-processing

Optionally, scReadSim allows users to design ground-truth open chromatin regions and then generates synthetic scATAC-seq reads accordingly. When users take this option, scReadSim requires users to input the BAM file, the trustworthy peaks and non-peaks of the BAM file, and the user-designed ground-truth peaks (Fig. S16). Given the user-designed ground-truth peaks, scReadSim specifies the complementary regions to the ground-truth peaks as the ground-truth non-peaks. In summary, scReadSim defines two sets of peaks and non-peaks:

(1) the set of “trustworthy peaks and non-peaks” based on the user-specified (or scReadSim-defined) trustworthy peaks and non-peaks and (2) the set of “ground-truth peaks and non-peaks” based on the user-designed ground-truth peaks.

For the “trustworthy peaks and non-peaks” feature set, scReadSim counts the number of reads overlapping each trustworthy peak or non-peak in every cell to construct a trustworthy-peak-by-cell count matrix and a trustworthy-non-peak-by-cell count matrix. The two matrices contain the same cells in the same column order.

For the “ground-truth peaks and non-peaks” feature set, since the specified ground-truth peaks may not correspond to real peaks in the BAM file, scReadSim constructs the ground-truth peaks’ counts by mapping the trustworthy peaks to the “most similar” ground-truth peaks in terms of region length. Below are the detailed construction steps.

(a) For every ground-truth peak, scReadSim defines a corresponding trustworthy peak as follows. First, scReadSim finds the ground-truth peak’s 50 most similar trustworthy peaks (in the same chromosome) in terms of lengths (Fig. S17a). Second, scReadSim chooses the trustworthy peak with the largest ratio of read count by peak length.

(b) To construct the ground-truth-peak-by-cell count matrix, for every ground-truth peak, scReadSim adds the corresponding trustworthy peak’s row in the trustworthy-peak-by-cell count matrix to the ground-truth-peak-by-cell count matrix (Fig. S17b). This constructed ground-truth-peak-by-cell count matrix has the same column order as the trustworthy-peak-by-cell count matrix.

The ground-truth-non-peak-by-cell matrix is constructed similarly, where every ground-truth non-peak is mapped to a trustworthy non-peak. Note that for each ground-truth non-peak, scReadSim first finds the 50 most similar trustworthy non-peaks in terms of lengths, and then it chooses the trustworthy non-peak with the smallest ratio of read count by peak length.

The ground-truth-peak-by-cell and ground-truth-non-peak-by-cell count matrices, as well as the maps from ground-truth features to trustworthy features, will be used in the next steps.

2. Synthetic count matrix generation

Similar to the synthetic count matrix generation for scATAC-seq without user-designed ground-truth peaks, scReadSim trains scDesign2 on the ground-truth-peak-by-cell and ground-truth-non-peak-by-cell count matrices (constructed from step “1. Pre-processing”) to generate the ground-truth-peak-by-cell and ground-truth-non-peak-by-cell count matrices for synthetic cells.

3. Synthetic read generation

To generate synthetic scATAC-seq reads from the synthetic ground-truth-peak-by-cell and ground-truth-non-peak-by-cell count matrices, for each ground-truth feature (peak or non-peak) in every synthetic cell, scReadSim (1) extracts the corresponding count *c* (i.e., the number of synthetic reads) from the synthetic ground-truth-feature-by-cell count matrices;

(2) samples (with replacement) *Ic/*2*1* real reads in the BAM file that overlap the ground-truth feature’s corresponding trustworthy feature; (3) obtain the mates of the *Ic/*2*1* sampled real reads; and (4) generates *Ic/*2*1* synthetic read pairs from the *Ic/*2*1* real read pairs, the read length, the signed distance between the ground-truth feature and its corresponding real feature, and the reference genome.

Specifically, in (4), scReadSim converts the *c* real reads’ 5’ positions in the reference genome to the *c* synthetic reads’ 5’ positions after two adjustments: first, shifting real reads from the trustworthy feature to the ground-truth feature based on the signed distance between the two features; second, adding a random shift to each real read pair’s 5’ positions. In greater detail, in the first adjustment step, assuming that the signed distance between the trustworthy and ground-truth features is +*d*, if a real read has a 5’ position *x*, then the synthetic read’s 5’ position is *x* + *d*. This adjustment ensures that the synthetic reads are located in the ground-truth feature as how the real reads are located in the real feature. The second adjustment (random shift addition) is to avoid duplicate reads due to sampling with replacement. Specifically, scReadSim adds the same random shift to a pair of reads’ 5’ positions to keep the fragment size unchanged after the shift.

### Output in FASTQ or BAM formats

To output the synthetic reads (excluding the UMI-based scRNA-seq reads 1, which contain cell barcodes and UMIs) in a FASTQ or BAM file, scReadSim first records the coordinates of the synthetic reads in a BED file. Then scReadSim uses the bedtools [31] getfasta function to extract every synthetic read’s sequence from the reference genome and store the synthetic reads’ sequences in a FASTA file, which is then converted to a FASTQ file by the seqtk software [32]. For paired-end reads, scReadSim would generate two FASTQ files: one for reads 1 and the other for reads 2. Users may choose FASTQ as the output format of scReadSim, or they may additionally output a BAM file by aligning the synthetic reads to the reference genome by the alignment software bowtie2 [33].

### Substitution errors introduced into synthetic reads

To mimic the sequencing error rates observed in the actual Illumina sequencing reads, scReadSim introduces substitution errors into the synthetic reads. Specifically, scReadSim first uses the software fgbio [34] to calculate every position *i*’s average error rate (i.e., the probability that the position is sequenced as a wrong nucleotide, denoted by *p_i_*) and three substitution error rates (i.e., the probabilities that the position is wrongly sequenced as the three other nucleotides; the sum of the three probabilities is *p_i_*) based on all real reads mapped to that position. Then for each synthetic read, scReadSim decides if every base call, say at position *i*, is erroneous by sampling from Bernoulli (*p_i_*). If the base call is decided to be erroneous (i.e., the Bernoulli sample is 1), a substitution nucleotide is randomly sampled from the other three nucleotides with probabilities proportional to the three substitution error rates at position *i*. Specifically, for paired-end sequencing technologies like scATAC-seq, scReadSim introduces substitution errors separately for the two mates in a pair.

## Data analysis

### Count-level comparison between synthetic and real data

To verify that scReadSim generates synthetic data that mimics the real data regarding the read-count level, we compared synthetic and real gene-by-cell UMI count matrices through the following aspects: summary statistics, correlations among top-expressed genes (or peaks), and visualization of two-dimensional (2D) embeddings. For scATAC-seq, in addition to these three aspects used in scRNA-seq, we compared the RPKM distributions between peak-by-cell read count matrices.

- **Summary statistics**. We utilized the following summary statistics to describe the marginal distributions of the count matrix entries: for every feature, we calculated the mean, variance, coefficient of variance, and zero proportion across cells; for every cell, we calculated the library size and zero proportion across features.
- **Feature-feature correlations**. To illustrate that scReadSim preserves the correlations among the genes (or peaks), we selected 100 top-expressed features (genes for scRNA-seq; peaks for scATAC-seq) from the real count matrix and used a heatmap to show the correlations among these features for both real and synthetic count matrices.
- **Visualization of 2D embeddings.** We utilized principal component analysis (PCA) and uniform manifold approximation and projection (UMAP) [35, 36] to obtain the 2D embedding of the combined real and synthetic count matrices: we first used PCA to obtain the 30 principal components (PCs) and then deployed UMAP to obtain the 2D embeddings. We labeled the cells with the dataset sources (synthetic cells generated by scReadSim or real cells). We also reported the median of the integration local inverse Simpson’s index (miLISI) proposed by [37] to quantify the mixing levels of synthetic and real cells (see Subsection ”Evaluation metrics”). The miLISI value (ranging from 1 to 2) will be close to 2 if a perfect mixture exists between the synthetic and real cells.
- **RPKM**. We used the quantity RPKM (see Subsection ”Evaluation metrics”) to summarize the read coverage over the whole reference genome. Specifically, we reported the RPKM quantities separately for the peaks and non-peaks.

### Sequence-level comparison between synthetic and real scRNA-seq data

We deployed scReadSim onto the mouse 10x single-cell Multiome dataset [20] (the RNA-seq modality only) by inputting the reference genome, the BAM file, and the gene annotation GTF file (Fig. 1a). Lastly, scReadSim generated synthetic scRNA-seq reads in the BAM format.

To show that the scReadSim’s synthetic scRNA-seq data mimics real data under the read-sequence level, we compared the real and synthetic BAM files through the k-mer spectrum, error rate per base, and genome browser visualization.

- **K-mer spectrum**. K-mer spectrum measures the occurrences of substrings of length *k* within the read sequences. We used the tool Jellyfish [38] with the default settings to obtain the 11-mer, 21-mer, 31-mer, and 41-mer spectra in the synthetic and real BAM files.
- **Error rates per base**. We calculated and compared the substitution error rate for every base in the real and synthetic reads. The substitution error rate per base within reads was obtained using the software fgbio [34] with the function ErrorRateByReadPosition by setting the option collapse as false.
- **Genome browser visualization**. We used the IGV genome browser [25] to visualize the real and synthetic BAM files. This visualization depicts how the pooled single-cell reads align with the reference genome and illustrates the bulk-level similarity of read coverage between the two BAM files.

### Sequence-level comparison between synthetic and real scATAC-seq data

We deployed scReadSim onto the sci-ATAC-seq and mouse 10x single-cell Multiome dataset [20] (the ATAC-seq modality only) by inputting the reference genome, the BAM file, and trustworthy peaks and non-peaks (Fig. 1b). The trustworthy peaks were identified using MACS3 from the input BAM file with a stringent rule (by setting q-value parameter as -q 0.01). To obtain the trustworthy non-peaks, we first deployed MACS3 onto the input BAM file with a less stringent rule (by setting q-value parameter as -q 0.1), then we took the inter-peaks as the trustworthy non-peaks. Lastly, scReadSim generated synthetic scATAC-seq reads in the BAM format.

To justify that scReadSim simulates realistic scATAC-seq data under the read-sequence level, we compared the real and synthetic BAM files through the k-mer spectrum, error rate per base, genome browser visualization (see Subsection ”Sequence-level comparison between synthetic and real scRNA-seq data”), and, additionally, the fragment size distribution. We further verified the realism of scATAC-seq data by deploying the peak caller MACS3 onto both real and synthetic BAM files and comparing the peak calling consistency.

- **Fragment-size distribution**. For paired-end sequencing reads, the fragment-size distribution measures the distribution of the distances between two ends of the read pairs. We used the function bamPEFragmentSize of the software deeptools [39] with the default settings to obtain the fragment distributions of real and synthetic scATAC-seq reads.
- **Peak calling comparison**. We deployed the peak-calling tool MACS3 with the same setting (by setting q-value parameter as -q 0.01) onto the real and synthetic BAM files and compared the peak calling results using a Venn diagram. In addition, to show the ROC (receiver operating characteristic) curve and Precision-Recall curve, we implemented MACS3 onto synthetic reads with a less stringent criterion to obtain more synthetic peaks for comparison (by setting q-value parameter as -q 0.5). Treating the peaks called from real data as truth, each peak identified from synthetic data is determined to be a true peak if it overlaps with a true peak over more than half of the minimal true peaks’ length. Specifically, we gradually changed the thresholds (such as p-values of peaks outputted by MACS3) of peak-calling tools to vary the number of peaks identified by the tools.

### Benchmark of UMI deduplication tools

We deployed scReadSim to the mouse 10x single-cell Multiome dataset [20] (the RNA-seq modality only) to generate the synthetic scRNA-seq data. Then we used scReadSim’s synthetic scRNA-seq reads as the input data and scReadSim’s gene-by-cell UMI-count matrix as the ground truth to benchmark three UMI deduplication tools: UMI-tools and cellranger (gene-level), as well as Alevin (transcript-level). We applied each deduplication tool to scReadSim’s synthetic scRNA-seq reads to obtain a deduplicated UMI count matrix. Since these deduplication tools may output different numbers of cells, we only considered the common cells outputted by all the tools for a fair comparison. We also removed the genes with zero expression levels in all cells in any deduplicated UMI count matrices or the ground-truth UMI count matrix. We compared the deduplicated UMI count matrices with the ground-truth count matrix in the following three aspects.

1. We calculated associations between the ground-truth UMI count matrix and each deduplicated UMI count matrix, gene-wise and cell-wise. The association measures include the Pearson correlation and Kendall’s tau.

2. We compared the UMI count matrices using the distributions of six summary statistics, including four gene-level statistics (mean, variance, coefficient of variance, and zero proportion) and two cell-level statistics (zero proportion and cell library size).

3. We used PCA and UMAP for dimension reduction and visualization to examine whether the deduplicated UMI count matrices preserve the 2D cell embeddings of the ground-truth UMI count matrix. Specifically, we first log-transformed the three UMI count matrices. Then we performed PCA on the ground-truth matrix to obtain the top 30 PCs and the associated 30 loading vectors. Next, we applied UMAP to the top 30 PCs to find the 2D cell embeddings. For the deduplicated matrices, we first projected their cells to the same 30-dimensional PC space by left multiplying the 30-by-*p* loading matrix (with rows as the top 30 loading vectors; *p* is the number of genes); second, we projected the deduplicated matrices’ cells from the PC space to the same UMAP space using the predict() function in the R package umap (Version 0.2.7.0).

In addition, we measured the time complexity by deploying each deduplication tool to scReadSim’s synthetic data with varying cell numbers: 1219 (0.25*×*), 2438 (0.5*×*), 4877 (1*×*, the original cell number), 9754 (2*×*), 19508 (4*×*); and varying sequencing depth 1.2*M* (0.25*×*), 2.5*M* (0.5*×*), 5.0*M* (1*×*, the original sequencing depth), 10.1*M* (2*×*), 20.1*M* (4*×*). The analysis was implemented on a server with the 256x Intel® Xeon Phi^TM^ CPU 7210 at 1.30GHz.

### Benchmark of peak-calling tools

To generate synthetic scATAC-seq reads with specified ground-truth peaks (i.e., open chromatin regions), we provided scReadSim with the mouse sci-ATAC-seq BAM file [22], the trustworthy peaks and non-peaks called from the BAM file (see Subsection “Sequence-level comparison between synthetic and real scATAC-seq data”), and a list of specified ground-truth peaks (Fig. S16). We specified each ground-truth peak as [TSS, TSS+LEN*−*1], where TSS is a gene’s transcription starting site, and LEN is uniformly sampled from [250, 550] bp. Then we merged any overlapping ground-truth peaks so that the final ones were non-overlapping. Taking these ground-truth peaks as input, scReadSim generated synthetic scATAC-seq reads in a BAM file.

To benchmark the peak-calling tools, MACS3, HOMER, SEACR, and HMMRATAC, we deployed these four peak-calling tools with the default settings onto the synthetic scATAC-seq BAM file with the specified ground-truth peaks. Specifically, MACS3 was run under -BAMPE mode with q-value 0.05; HOMER was run under mode -region provided parameter -minDist 150; SEACR was run with control threshold level 0.5 under -stringent mode; HMMRATAC was input required files with default settings. Then we compared the peak-calling results with the ground-truth peaks, throughout the following three aspects:

- First, we calculated the synthetic reads’ RPKM quantities for the ground-truth peaks and the peaks identified by the four peak-calling tools. The RPKM quantities are also reported for the non-peaks. We calculated the mean differences of peaks’ RPKMs between the ground truth and each method.
- Second, treating the ground-truth peaks as truth, we plotted the ROC and precision-recall curves by changing the number of discoveries for each peak-calling tool. A called peak is considered true if it overlaps at least 225 bp of a designed open chromatin region, where 225 bp is half of the shortest open chromatin region.
- Third, we directly compared these peak calling results through the Venn diagram and upset plot using the Python package Intervene [40].

### Comparison of scReadSim with minnow

We first deployed scReadSim to the mouse 10x single-cell Multiome dataset [20] (the RNA modality only) to generate synthetic reads in a BAM file. Since minnow does not accept reads but only a gene-by-cell UMI count matrix as input, we inputted into minnow the synthetic gene-by-cell UMI count matrix (generated by scReadSim as an intermediate step) and kept minnow’s parameters as default. The synthetic reads minnow generates were outputted in FASTQ files, and we aligned them to the same reference genome scReadSim uses. After binning the reference genome (chromosome 1) into 1,000-bp windows, we plotted the read coverage per window for the real data, scReadSim’s synthetic data, and minnow’s synthetic data (Fig. 1c). We further calculated the correlations (Spearman correlation and Pearson correlation) of read coverage between real and synthetic data across the windows. We also provided a closer view of the read coverage (region chr1:86,425,858–86,443,378) using the IGV genome browser (Fig. 1c Inset).

### Comparison of scReadSim with SCAN-ATAC-Sim

We used the sci-ATAC-seq dataset [22] to train scReadSim and SCAN-ATAC-Sim. Then we compared the two simulators’ performance in preserving the peak-count information in real data.

For scReadSim, we inputted the real dataset’s BAM file, the reference genome, and the trustworthy peaks (called at q-value 0.01) and non-peaks (complementary to the peaks called at q-value 0.1) identified by MACS3 from the BAM file. For each cell type, we used scReadSim to generate the same number of synthetic cells as the number of real cells. Moreover, we recorded the trustworthy peaks for the peak-count analysis.

Since SCAN-ATAC-Sim requires per-cell-type input, we focused on the four largest cell types with the most cells (predefined in [22], including Hematopoietic progenitors, Erythroblasts, Monocytes, and Immature B cells). Hence, we only examined the real reads and scReadSim’s synthetic reads for the four cell types.

For SCAN-ATAC-Sim to generate scATAC-seq data for the four cell types, it requires the four cell types’ pseudo-bulk ATAC-seq BAM files and their corresponding peak lists as input. Hence, we first extracted the four cell types’ reads from the real dataset’s BAM file to construct the four corresponding per-cell-type BAM files. Then we used MACS3 to call peaks from each per-cell-type BAM file. Finally, we inputted the four per-cell-type peak lists along with the four per-cell-type BAM files into SCAN-ATAC-Sim. For SCAN-ATAC-Sim’s parameters, we specified the cell number of each cell type to the real cell number, which is also scReadSim’s synthetic cell number, of the cell type. For other parameters, we used the default values suggested in the SCAN-ATAC-Sim paper [19]. SCAN-ATAC-Sim does not output synthetic read sequences but only a BED file containing synthetic reads’ starting and ending positions, along with the reads’ cell IDs.

To evaluate scReadSim and SCAN-ATAC-Sim’s performance at the peak-count level, we constructed scReadSim and SCAN-ATAC-Sim’s peak-by-cell read count matrices (one per simulator) from scReadSim’s output BAM file and SCAN-ATAC-Sim’s output BED file, by counting each simulator’s synthetic reads that overlap with the trustworthy peaks in the synthetic cells. We also constructed a real count matrix in the same way, so the three count matrices have the same dimensions.

In three aspects, we compared the three peak-by-cell count matrices (real, scReadSim, and SCAN-ATAC-Sim). First, we calculated the distributions of six summary statistics based on the entries of each count matrix: peak-wise mean, variance, coefficient of variance, and zero proportion across cells, as well as cell-wise library size and zero proportion across peaks. We further performed the two-sample Kolmogorov–Smirnov (KS) test to compare each summary statistic’s distributions in the real count matrix and a synthetic count matrix (scReadSim or SCAN-ATAC-Sim) to measure the resemblance of the synthetic count matrix to the real count matrix (a smaller KS statistic value indicates greater similarity).

Second, we obtained the 2D UMAP cell embeddings of the real and synthetic count matrices using the following steps.

1. We transformed the count matrices by applying the Term Frequency–Inverse Document Frequency (TF-IDF) transformation followed by the logarithmic transformation.

2. We performed PCA on the real pre-processed matrix to obtain the top 30 PCs and the associated 30 loading vectors. Next, we applied UMAP to the top 30 PCs to find the 2D cell embeddings.

3. For the synthetic pre-processed matrices, we projected their cells to the same 30-dimensional PC space by left multiplying the 30-by-*p* loading matrix (with rows as the top 30 loading vectors; *p* is the number of peaks). Then we projected the synthetic cells from the PC space to the same UMAP space using the predict() function in the R package umap (Version 0.2.7.0). We labeled the cells with the cell types. We reported the miLISI (proposed by [37]) to quantify the mixing levels of synthetic and real cells (see Subsection “Evaluation metrics”). The miLISI value (ranging from 1 to 2) will be close to 2 if a perfect mixture exists between the synthetic and real cells.

Third, we visualized how scReadSim’s and SCAN-ATAC-Sim’s synthetic cells mix with real cells in the 2D UMAP embeddings using the following steps.

1. We combined real cells and scReadSim’s synthetic cells by concatenating the real peak-by-cell pre-processed matrix and scReadSim’s synthetic peak-by-cell pre-processed matrix (from TF-IDF followed by the logarithmic transformation) horizontally as a peak-by-mixed-cell matrix. Similarly, we combined real cells and SCAN-ATAC-Sim’s synthetic cells into a concatenated peak-by-mixed-cell matrix.

2. We projected the two groups of mixed cells to the previously used 30-dimensional PC space by left multiplying the 30-by-*p* loading matrix (with rows as the top 30 loading vectors; *p* is the number of peaks) obtained previously from the real pre-processed matrix.

3. We projected the two groups of mixed cells from the PC space to the previously used UMAP space using the predict() function in the R package umap (Version 0.2.7.0).

### Evaluation metrics

#### Reads Per Kilobase Million (RPKM)

The RPKM value of a feature is defined as

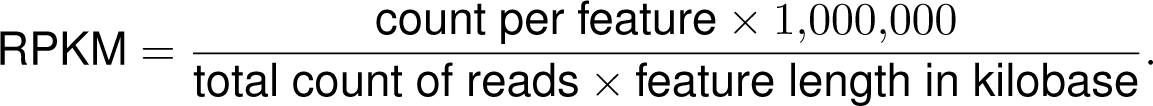

#### miLISI

The integration local inverse Simpson’s Index (iLISI) [37] for each cell measures the effective number of datasets in a neighborhood of the cell. A cell surrounded by neighboring cells from a single dataset has an iLISI value equal to 1; otherwise, if the cell has a neighborhood with an equal number of cells from two datasets, the iLISI value would be 2. Hence, the median iLISI (miLISI) across all cells measures the median mixing level of the two datasets, and a miLISI close to 2 means that the two datasets have a perfect mixing.

## Notes

https://screadsim.readthedocs.io/

### Competing Interest Statement

The authors have declared no competing interest.

### Summary of Updates

Main figure revised; authorship updated.

